# TMEM106B mediates ACE2-independent replication of the SARS-CoV-2 S^E484D^ variant in airway-derived cell models

**DOI:** 10.64898/2026.03.14.711762

**Authors:** Ming Xia, Qibo Wang, Maritza Puray-Chavez, Kyle LaPak, Zhuoming Liu, Sathvik Palakurty, Paul Rothlauf, Sean Hui, Natalie Stegman, Marjorie Cornejo Pontelli, Jacob Marsh, Anayat Bhat, Celeste M. Karch, Daved H. Fremont, Ramon Lorenzo-Redondo, Michael S. Diamond, Sean P.J. Whelan, Michael B. Major, Dennis Goldfarb, Sebla B. Kutluay

**Author notes:** Address correspondence to Sebla B. Kutluay.

## Abstract

Severe acute respiratory syndrome coronavirus 2 (SARS-CoV-2) continues to cause significant respiratory disease, particularly in vulnerable populations. Although ACE2 is the primary receptor for viral entry, previous studies have identified a naturally occurring, ACE2-independent entry pathway that depends on cell surface heparan sulfates and the E484D substitution within the receptor-binding domain of Spike (S). In this study, we expand the panel of airway-derived cell lines that support ACE2-independent, S-E484D-dependent entry and identify the host lysosomal transmembrane protein TMEM106B as the alternative receptor for infection. Specifically, we show that knockout of TMEM106B completely abolishes infection by SARS-CoV-2 S^E484D^ in NCI-H522 and NCI-H661 cells. Ectopic expression of the luminal C-terminal domain (CTD) of TMEM106B-either alone or redirected to the plasma membrane – is sufficient to enable viral entry largely independent of cell surface heparan sulfates in non-permissive cells. We show that S-E484D is required for TMEM106B utilization, whereas other Variants of Concern substitutions at S-E484 are unable to mediate entry. Fc-TMEM106B-CTD fusion protein binds to a chimeric VSV-SARS-CoV-2 S^E484D^ with a higher affinity than WT virus and blocks infection by SARS-CoV-2 S^E484D^, supporting a direct interaction between S-E484D and TMEM106B-CTD. Serial passaging experiments with VSV-SARS-CoV-2 S^E484D^ identify a V951A mutation within the heptad repeat 1 region that enhances viral spread and fusogenicity in the ACE2-independent cell models. Together, these findings demonstrate that TMEM106B is a key mediator of a naturally occurring ACE2-independent pathway in multiple airway-derived cell lines and suggest that variation in the S protein can expand receptor usage by SARS-CoV-2.

**IMPORTANCE:** Severe acute respiratory syndrome coronavirus 2 (SARS-CoV-2) continues to acquire mutations in the viral spike (S) protein as it circulates in humans. How these mutations influence the cell and tissue tropism of SARS-CoV-2 remains poorly understood. While ACE2 is the canonical host cell receptor for SARS-CoV-2, previous studies revealed the presence of a naturally occurring ACE2-independent entry pathway in multiple airway-derived cell lines that can be utilized only by a SARS-CoV-2 variant bearing the E484D substitution within the viral S protein. Here we expand the airway-derived cell line models that support ACE2-independent and S-E484D-dependent entry and show that the host lysosomal transmembrane protein, TMEM106B, is an essential viral entry factor. Although the E484D substitution is currently rare in human isolates, our findings raise the possibility that even single mutations in SARS-CoV-2 S can allow the utilization of alternative entry pathways that broadens its cell and tissue tropism.

## INTRODUCTION

Severe acute respiratory syndrome coronavirus 2 (SARS-CoV-2) continues to cause respiratory disease, particularly in vulnerable populations. The emergence of viral variants that evade vaccine– and infection-induced immunity remains a major public health concern. These variants continue to evolve, in part through immune-evasive mutations within the viral spike (S) protein. SARS-CoV-2 pathogenesis is complex and involves multiple organ systems beyond the airways (1, 2), most notably the central nervous system (CNS) in long COVID. However, it remains unclear whether CNS and other extrapulmonary manifestations result from direct viral infection or from dysregulated immune responses. Beyond the cognate host cell receptor, ACE2, molecular mechanisms that define cell and tissue tropism of SARS-CoV-2 are incompletely understood. A greater understanding of cell intrinsic mechanisms that enhance or curb SARS-CoV-2 infection could explain differences in organ involvement, dissemination within and between hosts, and identify new opportunities for therapeutic intervention.

A prevailing view is that SARS-CoV-2 entry into the host cell requires viral homotrimeric S glycoprotein binding to the host cell angiotensin converting enzyme 2 (ACE2) receptor (3–6). Notwithstanding this point, endogenous ACE2 expression is surprisingly low in the respiratory tract (7, 8) and several other organ systems are impacted by SARS-CoV-2. We previously identified an ACE2-deficient airway epithelial cell line, NCI-H522 (hereafter H522), that nevertheless supports SARS-CoV-2 entry, replication and productive infection (9). This alternative entry pathway depends on surface expression of heparan sulfates, clathrin-mediated endocytosis, and requires the E484D substitution within the receptor-binding domain (RBD) of S (S^E484D^) (9). Whether there is a proteinaceous host cell receptor that mediates S^E484D^-dependent entry in this infection model is unclear.

Recent studies reported that the host lysosomal transmembrane protein, TMEM106B, can mediate ACE2-independent and S^E484D^-mediated entry in various cell lines (10–12). However, whether TMEM106B can directly mediate SARS-CoV-2-S^E484D^ binding and entry is unclear for several reasons. First, unlike ACE2, which localizes at the plasma membrane, TMEM106B predominantly resides in late endosomes/lysosomes (13). Whether TMEM106B plasma membrane localization is required for virus binding is not known. Second, how heparan sulfates and possibly lectins contribute to TMEM106B-mediated entry is not established. Third, SARS-CoV-2 S has a ∼1,000-fold lower binding affinity to TMEM106B than ACE2 (10). Whether this low affinity is sufficient to mediate entry is not known. Fourth, TMEM106B can function independently as a proviral factor through its established roles in lysosomal biology (14). As such, altering its expression levels may indirectly affect virus replication downstream from entry, without directly affecting virus binding.

Herein, we identified an additional airway-derived cell line, NCI-H661 (hereafter H661), that naturally supports ACE2-independent entry and similarly requires the S^E484D^ substitution for replication. We found that S^E484D^-mediated infection of both H522 and H661 cells depends on TMEM106B expression. Specifically, genetic knockout of *TMEM106B* in both H522 and H661 cells abolished binding and replication of SARS-CoV-2-S^E484D^. Complementation of TMEM106B knockout cells with full-length TMEM106B or the luminal C-terminal domain (CTD) alone restored virus binding and infection, whereas expression of the N-terminal domain (NTD) did not. Targeting TMEM106B-CTD to the plasma membrane restored infection in otherwise non-permissive cells, including those lacking cell surface heparan sulfates, demonstrating that the TMEM106B-CTD is sufficient to mediate virus binding and entry. Furthermore, an Fc-TMEM106B-CTD decoy protein bound to S^E484D^ at a higher affinity than wild-type (WT) S and inhibited replication of SARS-CoV-2-S^E484D^, establishing a direct interaction between S-E484D and TMEM106B-CTD. Using a chimeric VSV-SARS-CoV-2 S virus, we observed that while the S^E484D^ substitution was required for infection, virus spread was relatively inefficient in H522 and H661 cells. Serial passaging of VSV-SARS-CoV-2 S^E484D^ in H522 cells selected for variants with progressively increased infectivity, while retaining the S^E484D^ substitution. This enhanced viral spread was associated with the presence of a V951A substitution within the heptad repeat 1 (HR1) of S, which restored fusogenicity of VSV-SARS-CoV-2 S^E484D^ in both ACE2-and TMEM106B expressing cells. Finally, through in depth analyses of sequencing data sets, we show that the S-E484D substitution is mutationally accessible but rare in nature, suggesting that it is likely selected against. Together, these results demonstrate that TMEM106B is a key host determinant of an ACE2-independent entry pathway mediated by the S-E484D Spike variant and highlight that small modifications in Spike can alter receptor utilization by SARS-CoV-2.

## RESULTS

### H661 epithelial lung carcinoma cells support S^E484D^-dependent SARS-CoV-2 replication

A previous screen of lung and upper airway cell lines identified the H522 lung adenocarcinoma cell line that supports ACE2-independent SARS-CoV-2 replication and required the presence of S^E484D^ substitution (9). However, numerous ACE2 expressing cells were non-permissive to infection due to high basal activation of the cGAS-STING pathway and subsequent interferon (IFN) signaling (15), whereas the H522 cells expressed markedly lower levels of antiviral IFN-stimulated genes (**Fig. 1A**, see also (15)). To identify additional cell lines that support ACE2-independent entry, we analyzed publicly available RNA expression patterns and selected H661 (lung carcinoma), H1341 (small cell lung carcinoma), H841 (small cell lung carcinoma) and SNU-182 (hepatocellular carcinoma) cells based on their low ACE2 expression (**Fig. 1B**) and lack of basal innate immune activation signatures (**Fig. 1A, 1B**). Comparison of the cellular surface protein expression patterns revealed that H522, H661 and H841 cell lines were the most similar, whereas H1341 and SNU-182 cells were more distally related (**Fig. 1C**). RT-qPCR and immunoblot analyses showed that H1341 cells express relatively high levels of ACE2 approaching that observed in the permissive Vero E6 cells (**Fig. 1D, 1E**). SNU-182 cells expressed low but detectable levels of ACE2 mRNA and protein (**Fig. 1D, 1E**), comparable to levels in H1299 cells, which we previously showed support ACE2-dependent SARS-CoV-2 replication (15). In contrast, H661 and H841 cells lacked detectable ACE2 mRNA or protein (**Fig. 1D, 1E**). Infection experiments using SARS-CoV-2 bearing WT S, S-E484D, or S-E484K showed that H1341 and SNU-182 cells were readily infected by all variants, despite the notable ACE2 expression level differences **(Fig. 1F)**. In contrast, H661 cells were infected only by the S-E484D variant, and H841 cells did not support infection by any variant **(Fig. 1F)**. Together, these data show that like H522 cells, H661 cells require the presence of S-E484D Spike variant for infection.

**Figure 1.**
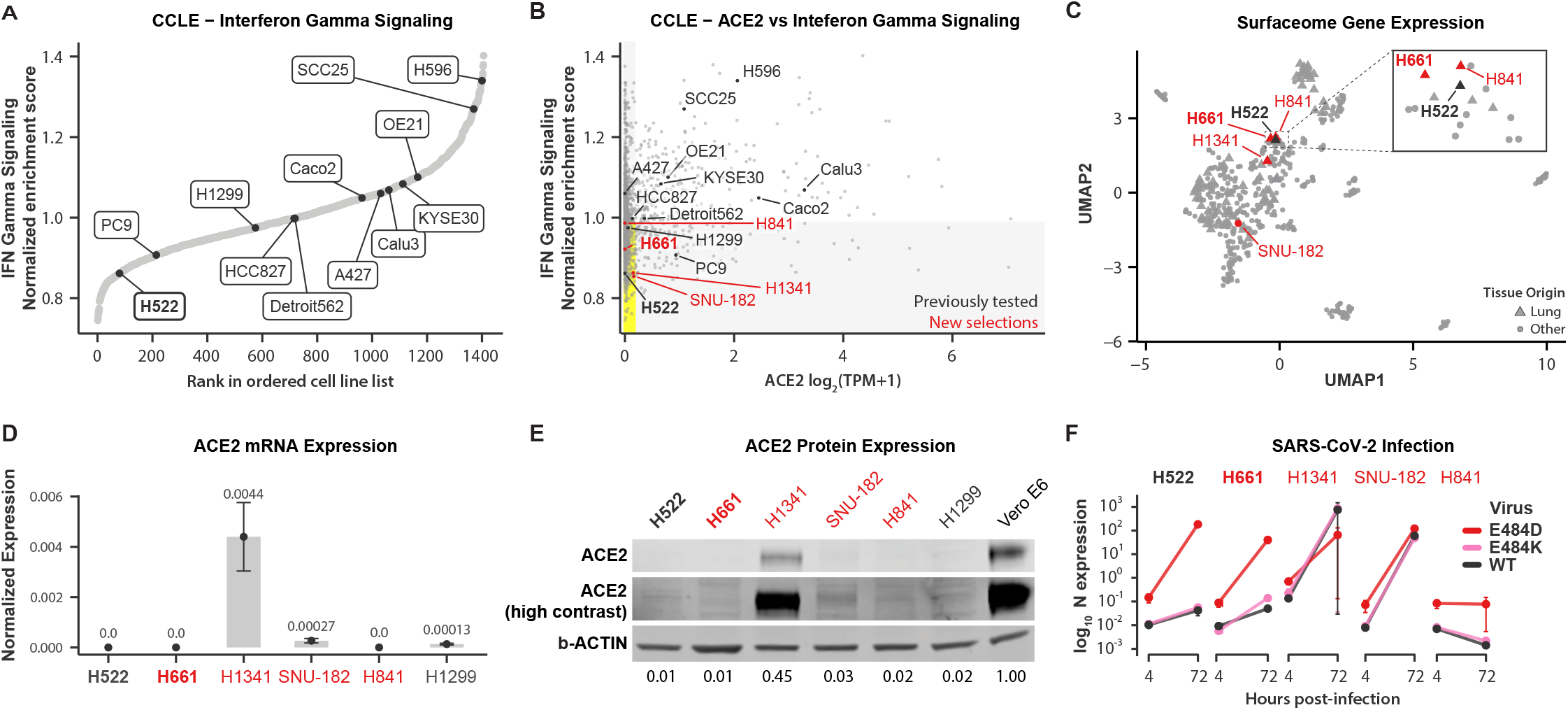
H661 lung adenocarcinoma cells are preferentially infected by the SARS-CoV-2 S^E484D^ variant. **(A)** Enrichment analysis of the expression levels of interferon gamma signaling genes in the indicated cell lines using publicly available RNA-seq data sets. **(B)** Comparison of the expression levels of ACE2 protein (x-axis) and mRNAs in the interferon gamma signaling pathway (y-axis) in the indicated cell lines. **(C)** UMAP analysis of cell surface protein expression in the indicated cell lines. **(D)** RT-qPCR analysis of ACE2 mRNA expression in the indicated cell lines. **(E)** Immunoblot analysis of ACE2 and β-actin expression in the indicated cell lines. **(F)** Cells were inoculated with SARS-CoV-2 (USA-WA1-2020) bearing WT S or S^E484D^ or B.1.526 SARS-CoV-2 variant bearing S-E484K substitution at an MOI of 1 i.u./cell. Cell-associated viral RNA levels was analyzed at 4 h and 72 h post-infection by RT-qPCR from n=3 independent replicates. Data show the mean, error bars show the SD.

### H661 cells support ACE2-independent SARS-CoV-2 replication

We next interrogated whether the H661 cells support ACE2-independent infection and whether this alternative entry pathway depends on endosome acidification and cell-surface heparan sulfates as seen before with H522 cells (9). Consistent with an ACE2-independent entry mechanism, vRNA levels in bulk populations of H661 ACE2 knockout (KO) cells were indistinguishable from those in WT cells at 4 and 72 hours post-infection (hpi), similar to observations in H522-ACE2 KO cells (**Fig. 2A**, also see (9)). Likewise, four independent H661-ACE2-KO monoclonal lines with Cas9-induced editing in the *ACE2* locus (**Fig. 2B**) supported robust replication comparable to parental controls (**Fig. 2C**), demonstrating that SARS-CoV-2 S^E484D^ infection of H661 cells occurs independently of ACE2. Treatment of both H522 and H661 cells with ammonium chloride, which neutralizes endosomal pH, markedly reduced cell-associated vRNA levels (Fig**. 2D**), consistent with a pH-dependent endosomal entry mechanism. In addition, enzymatic removal of cell-surface heparan sulfates reduced virus replication in both cell lines (**Fig. 2E**), supporting a model in which S^E484D^ utilizes heparan sulfate–mediated attachment prior to endosomal entry.

**Figure 2.**
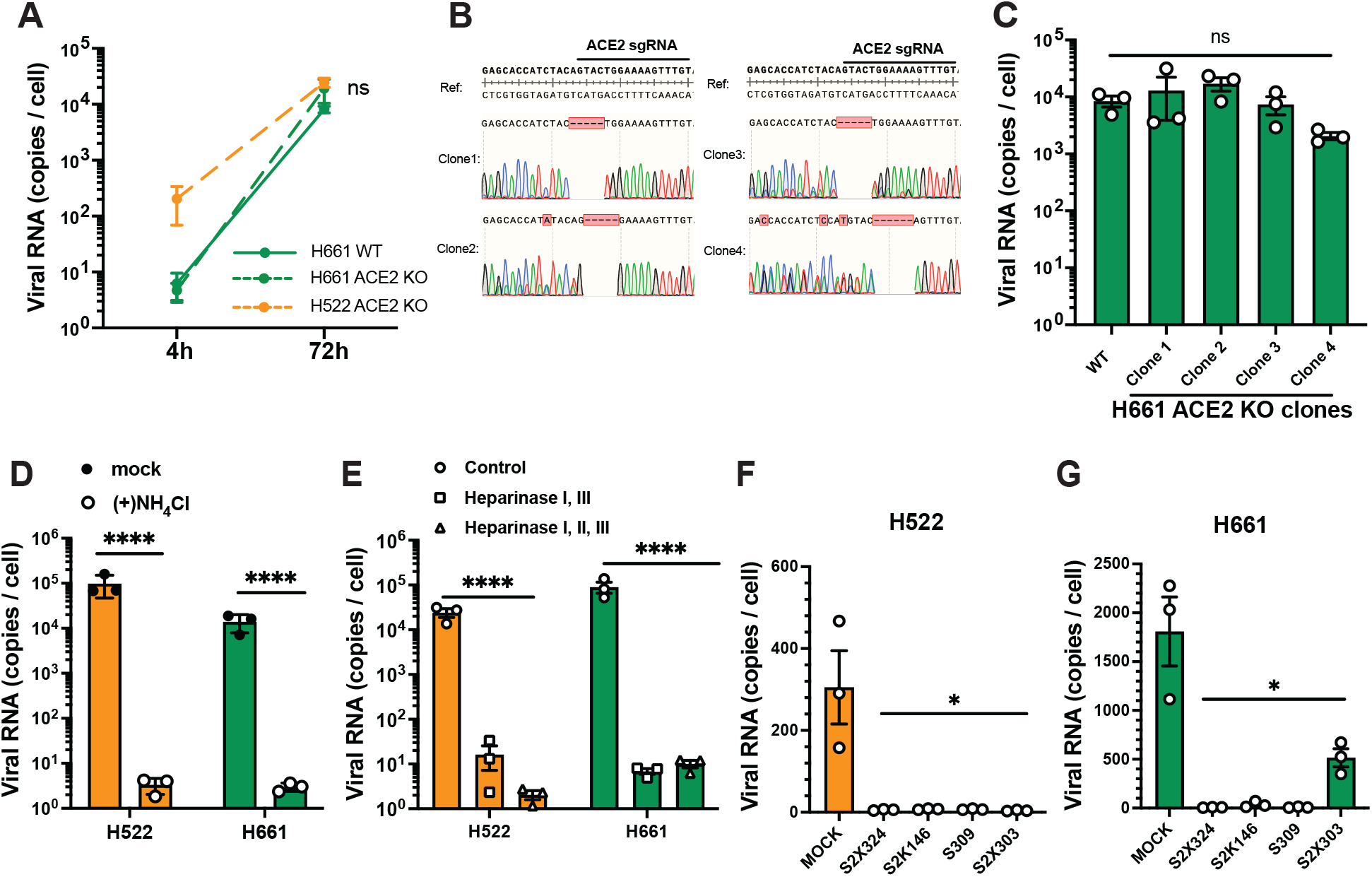
H522 and H661 cells support SARS-CoV-2 S^E484D^ replication independent of ACE2. **(A)** H522-ACE2 KO, H661 and H661-ACE2 KO cells were inoculated with SARS-CoV-2-mNG S^E484D^ at an MOI of 1. RT-qPCR analysis of cell-associated viral RNA at 4 and 72 hpi is shown. **(B)** Sanger sequencing of genomic ACE2 at exon 3. Data from unique monoclonal populations of H661 ACE2 KO cells were aligned to the human genome (‘Ref’; hg38). The dashed lines indicate small deletions within exon 3 of ACE2. **(C)** WT H661 and four different H661 ACE2 KO clonal cells were inoculated with SARS-CoV-2-mNG S^E484D^ at an MOI of 1. RT-qPCR for cell-associated viral RNA at 72 hpi is shown. **(D)** H522 and H661 WT cells pre-treated with 50 mM ammonium chloride for 2 h and inoculated with SARS-CoV-2-mNG S^E484D^ at an MOI of 1. At 24 hpi, ammonium chloride was removed, and cells were replenished with fresh media. RT-qPCR analysis of cell-associated SARS-CoV-2 RNA at 48 hpi is shown. **(E)** H522 and H661 cells were pre-treated with a combination of heparinase I, II and III for 90 min. Cells were subsequently inoculated with SARS-CoV-2-mNG S^E484D^ at an MOI of 0.1. RT-qPCR for cell-associated SARS-CoV-2 RNA at 72 hpi is shown. **(F, G)** H522 (E) and H661 cells (F) were infected with SARS-CoV-2-mNG S^E484D^ (MOI of 0.1) in the presence of mAbs. Amount of cell-associated vRNA was quantified as above. All data show the mean from three independent biological replicates with error bars displaying the SEM. (nonsignificant (ns); \*\**P* < 0.01; \*\*\**P* < 0.001; \*\*\*\**P* < 0.0001 by two-tailed unpaired *t*-test or one-way ANOVA with Dunnett’s correction for multiple comparisons).

We next evaluated whether monoclonal antibodies (mAbs) targeting distinct epitopes on the S protein could inhibit ACE2-independent infection mediated by the S^E484D^ variant. The broadly neutralizing antibodies S309, S2X324 and S2K146, which recognize conserved epitopes within the RBD (16), potently inhibited infection of SARS-CoV-2 S^E484D^ in both H522 and H661 cells (**Fig. 2F, 2G**). S2X303, which recognizes the N-terminal domain (NTD) of S (17) also blocked replication in H522 cells but had less neutralizing activity in H661 cells (**Fig. 2F, 2G**). These findings suggest that antibodies capable of engaging conserved RBD or NTD epitopes can block the ACE2-independent entry mechanism in H522 and H661 cells.

### TMEM106B is essential for S-E484D infection in ACE2-deficient cells

TMEM106B is a lysosomal transmembrane protein recently implicated in ACE2-independent entry (10, 11). We evaluated whether expression of TMEM106B is required for S^E484D^ infection in H522 and H661 cells. CRISPR-Cas9 knockout of TMEM106B was efficient, as validated by immunoblotting (**Fig. 3A**). TMEM106B KO resulted in a reduction in cell-associated SARS-CoV-2 RNA by ∼100-fold in H522 cells compared to parental and non-targeting (NT) controls (**Fig. 3B**) and led to a marked decrease in the number of infected cells (**Fig. 3C, 3D**). A similar requirement was observed in H661 cells: KO of TMEM106B expression (**Fig. 3E**) decreased cell-associated viral RNA levels (**Fig. 3F**) and reduced infection frequencies (**Fig. 3G, 3H**). Together, these data show that TMEM106B is required for SARS-CoV-2 S^E484D^ infection of H522 and H661 cells.

**Figure 3.**
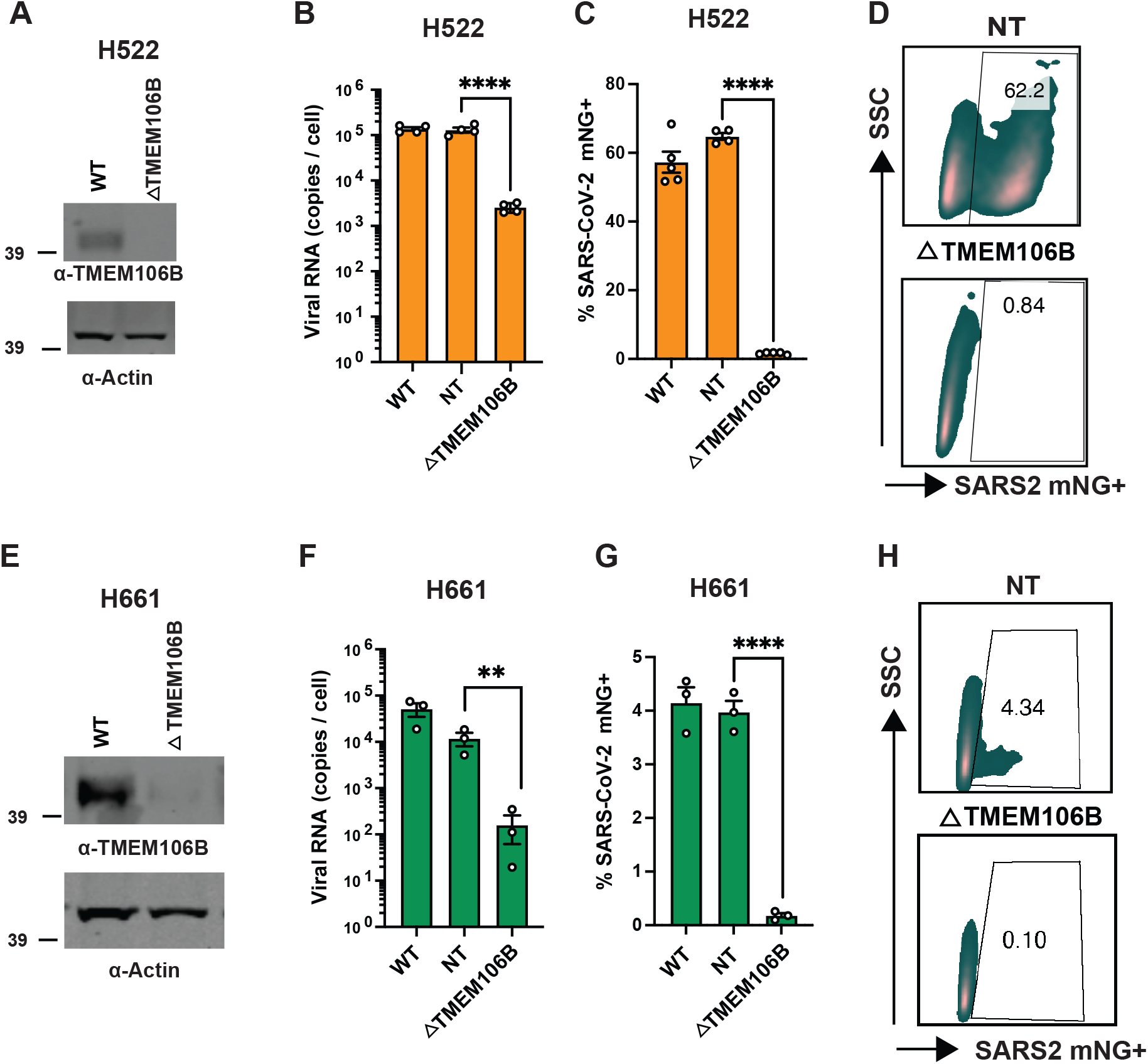
Knockout of TMEM106B abolishes infection of H522 and H661 cells. (**A-H**) H522 and H661 TMEM106B KO and non-targeting (NT) guide RNA-transduced cells were generated as detailed in Materials and Methods. TMEM106B expression in bulk populations of H522 (A) and H661 (E) cells was analyzed by immunoblotting using anti-TMEM106B and anti-actin antibodies. Parental wild-type (WT), TMEM106B KO and NT cells were inoculated with SARS-CoV-2-mNG S^E484D^ at an MOI of 1. RT-qPCR for cell-associated SARS-CoV-2 RNA (**B, F**) and enumeration of mNG positive cells by FACS (**C, D, G, H**) at 48 hpi is shown (\*\**P* < 0.01; \*\*\*\**P* < 0.001 by two-tailed unpaired *t*-test).

### The luminal CTD of TMEM106B is necessary and sufficient for infection of H522 and H661 cells

TMEM106B is a type II transmembrane protein that localizes primarily to late endosomes and lysosomes and comprises a cytosolic N-terminal domain (NTD), a transmembrane helix (TM), and a glycosylated luminal C-terminal domain (CTD) (**Fig. 4A**). To define the TMEM106B domain important for SARS-CoV-2 S^E484D^ infection, we performed complementation experiments in TMEM106B-KO H522 and H661 cells stably expressing HA-tagged TMEM106B variants (**Fig. 4A**). Expression levels of all HA-tagged constructs were verified by immunoblotting (**Fig. 4B, 4C**). Expression of full-length (FL) TMEM106B restored SARS-CoV-2 S^E484D^ replication to near-WT levels in H522 cells, whereas TMEM106B-ΔCTD failed to rescue (**Fig. 4D**). In contrast, TMEM106B-ΔNTD restored replication at levels comparable to FL-TMEM106B (**Fig. 4D**), indicating that the TMEM106B-CTD is required for SARS-CoV-2 S^E484D^ infection of H522 cells. These results were confirmed with the VSV-SARS-CoV-2 S^E484D^ chimeric virus, which similarly required the TMEM106B-CTD for infection (**Fig. 4E**). Similar domain requirements were observed in H661 cells: TMEM106B-FL and TMEM106B-ΔNTD complementation restored infection, whereas ΔCTD did not for both SARS-CoV-2 S^E484D^ and VSV-SARS-CoV-2 S^E484D^ (**Fig. 4F, 4G**). Together, these findings show that the luminal CTD of TMEM106B is necessary for SARS-CoV-2 S^E484D^ infection, whereas the cytosolic N-terminal tail is dispensable.

**Figure 4.**
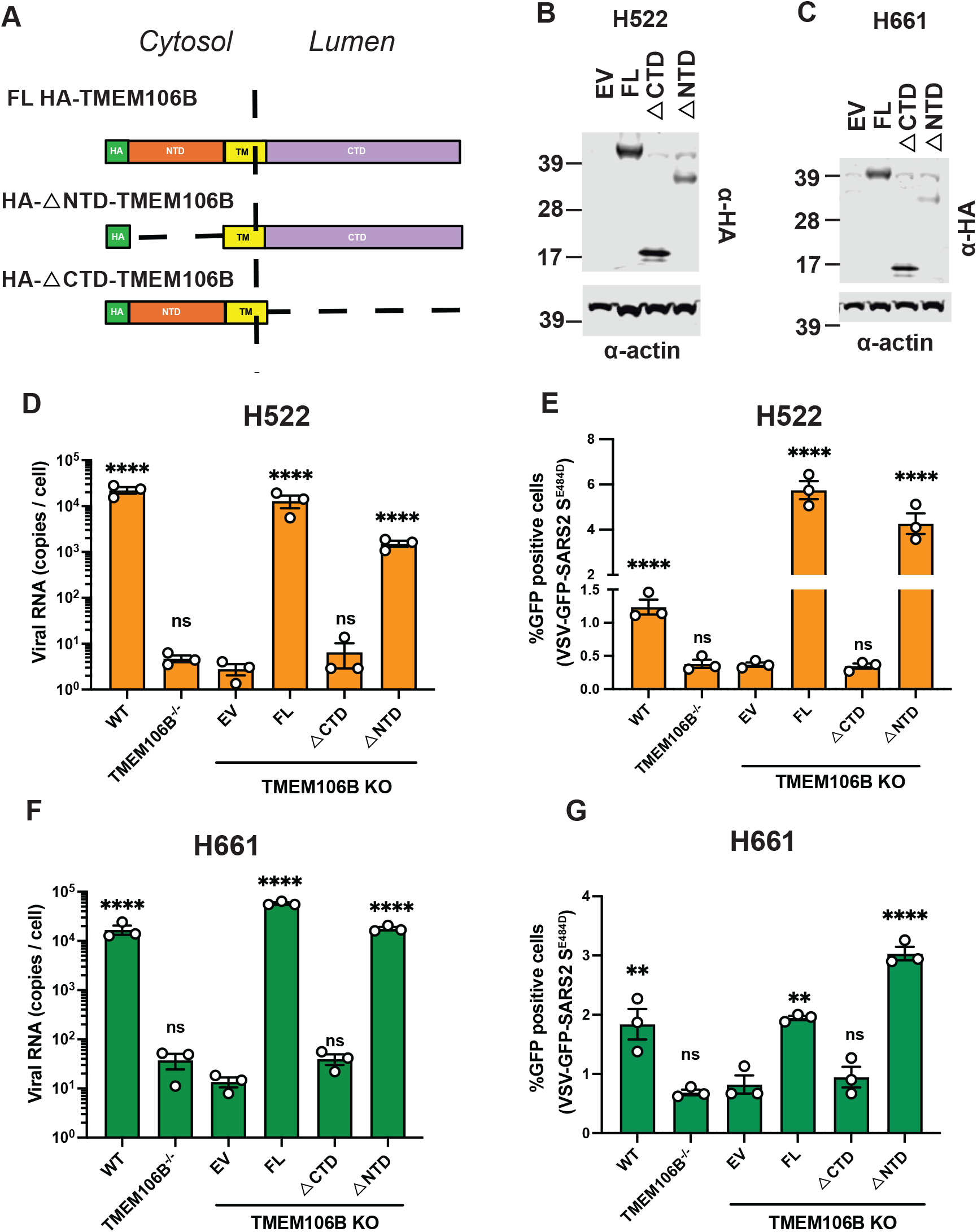
Expression of TMEM106B-CTD promotes SARS-CoV-2-S^E484D^ infection. **(A)** Schematic diagram of TMEM106B and deletion constructs used in this study. **(B-G)** TMEM106B KO H522 and H661 cells were stably transduced with an empty vector (EV), full-length (FL) TMEM106B, TMEM106BΔCTD, or TMEM106BΔNTD. Western blot analysis of H522 **(B)** and H661 **(C)** cell lysates using an anti-HA antibody is shown. Beta-actin was used as loading control. Indicated H522 and H661 cells were inoculated with SARS-CoV-2-mNG S^E484D^ (**D, F**) or VSV-GFP-SARS-CoV-2 S^E484D^ (**E, G**) at an MOI of 1. RT-qPCR analysis of cell-associated SARS-CoV-2 RNA at 72 hpi (**D, F**) and flow cytometric quantification for GFP positive cells at 48 hpi (**E, G**). Data in **D-G** show the mean of three independent biological replicates, error bars show the SEM. For statistical analysis, TMEM106B complemented cells were compared to EV-transduced cells. (non-significant (ns), \*\**P* < 0.01, \*\*\*\**P* < 0.0001 by one-way ANOVA with Dunnett’s correction for multiple comparisons).

### TMEM106B-CTD mediates virion binding and is sufficient for direct engagement of SARS-CoV-2 S^E484D^

To determine whether TMEM106B acts at the stage of virus binding and entry, we quantified SARS-CoV-2 S^E484D^ binding to H522 and H661 cells lacking TMEM106B or complemented with the aforementioned TMEM106B variants. Cells were inoculated with virus at 16°C to permit binding and were fixed immediately for RNAscope-based detection of incoming viral RNA. In both H522 and H661 TMEM106B-KO cells, virion binding was undetectable (**Fig. 5A**). Complementation with full-length TMEM106B measurably enhanced virus binding, and notably, expression of TMEM106B-ΔNTD at comparable levels (**Fig. 4B, 4C**) resulted in a substantially greater number of surface-associated vRNA puncta than in control cells (**Fig. 5A, 5B**). In contrast, TMEM106B-ΔCTD failed to enhance SARS-CoV-2 S^E484D^ binding (**Fig. 5A, 5B**). In agreement with the strict requirement for the S-E484D substitution in infection of H522 and H661 cells, WT SARS-CoV-2 did not bind to H522-TMEM106B-ΔNTD cells (**Fig. 5A, 5B**); instead, ectopic expression of ACE2 in H522 cells promoted WT SARS-CoV-2 binding (**Fig. 5A, 5B**). Finally, an Fc-TMEM106B-CTD soluble decoy protein blocked infection of H522 and H661 cells with SARS-CoV-2 S^E484D^ (**Fig. 5C, 5D**) and bound to VSV-GFP-SARS-CoV-2 S^E484D^ more efficiently than the WT virus (**Fig. 5E**) indicating that the luminal TMEM106B-CTD directly engages with S-E484D to mediate virion attachment.

**Figure 5.**
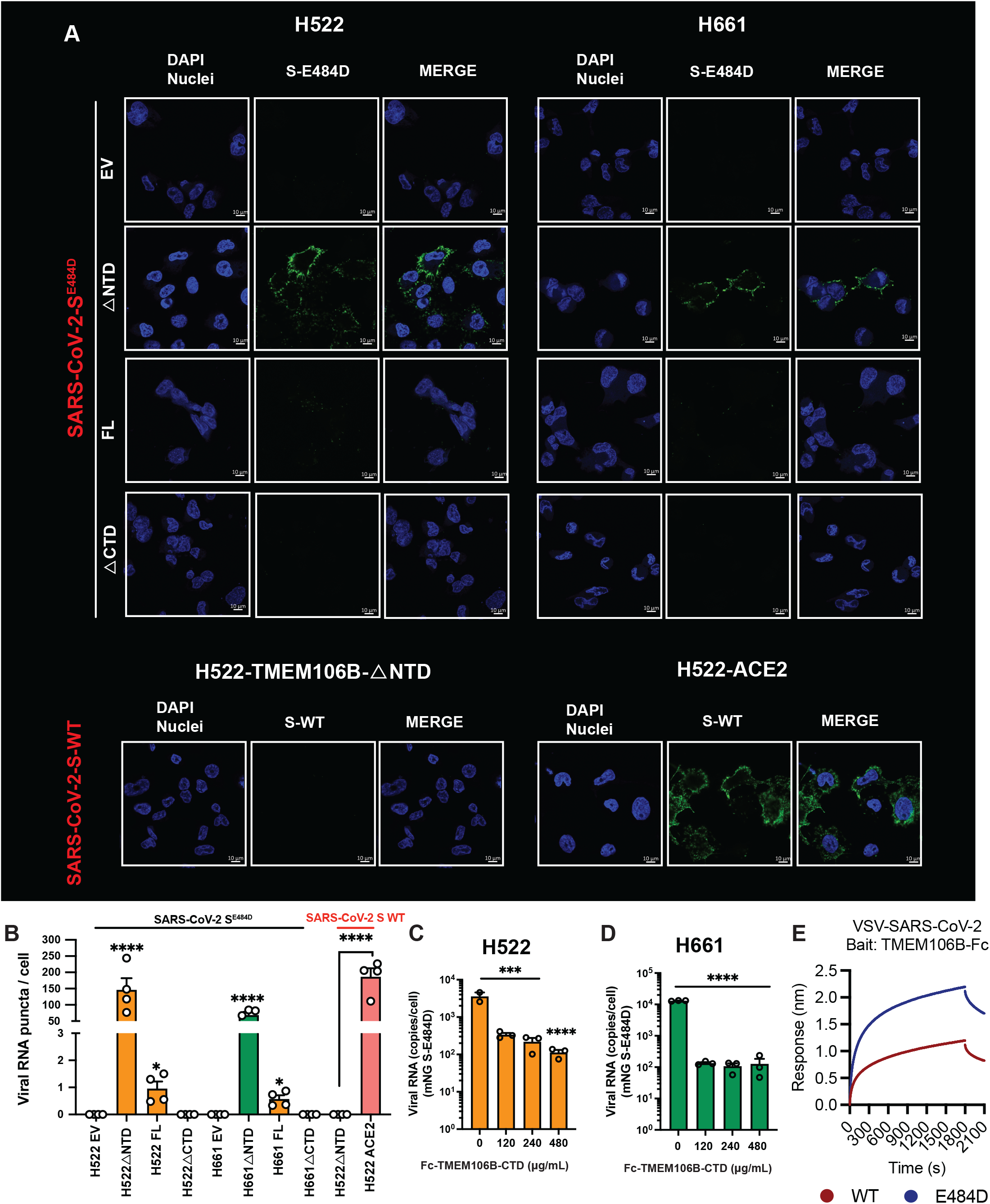
TMEM106B-CTD enhances binding of SARS-CoV-2-S^E484D^ but not the WT virus. (A) H522 and H661 TMEM106B KO cells stably transduced with an empty vector (EV), full length TMEM106B (FL), TMEM106BΔCTD, TMEM106BΔNTD as well as H522 cells stably expressing ACE2 were inoculated with SARS-CoV-2-mNG S^E484D^ or WT SARS-CoV-2-mNG at an MOI of 1 at 16°C. At 1 hpi, cells were fixed and analyzed for viral RNA (green) using RNAScope as detailed in Materials and Methods. Nuclear DAPI staining is shown in blue. (B) Quantification of the data shown in A. For S^E484D^ infections, all data was compared to EV, for S-WT infections H522-ACE2 OE data was compared to H522-TMEM106BΔNTD (\**P* < 0.05; \*\*\*\**P* < 0.0001 by two-tailed unpaired *t*-test or one-way ANOVA with Dunnett’s correction for multiple comparisons). Scale bar, 10 µm.μ (C, D) SARS-CoV-2 S^E484D^ (MOI of 2 equivalent) was pre-incubated with the indicated concentrations of Fc-TMEM106B-CTD and added on H522 (C) and H661 (D) cells as detailed in Materials and Methods. Cell-associated viral RNA was analyzed by RT-qPCR at 72 h post infection. Data show the average of three independent replicates with error bars displaying the SEM. \*\*\**P* < 0.001; \*\*\**P* < 0.0001 by two-tailed unpaired *t*-test or one-way ANOVA with Dunnett’s correction for multiple comparisons. (E) BLI sensogram of Fc-TMEM106B-CTD binding to VSV-SARS-CoV-2 S WT and S^E484D^.

### Targeting TMEM106B-CTD to the plasma membrane enhances SARS-CoV-2 S^E484D^ replication

Lysosomal localization of TMEM106B is mediated by an extended dileucine motif in its NTD (18). We speculated that the enhanced virus binding observed with TMEM106B-ΔNTD might be due to its enhanced plasma membrane localization. Consistent with this hypothesis, we found that while TMEM106B-FL and TMEM106B-ΔCTD colocalized with the late endosome/lysosome marker Rab7A, TMEM106B-ΔNTD preferentially localized to the plasma membrane, marked by WGA staining (**Fig. 6A**). We next tested whether the luminal CTD of TMEM106B is sufficient to support virus replication when redirected to the plasma membrane. To this end, we engineered a chimeric construct in which TMEM106B-CTD was fused to the N-terminal domain of the transmembrane protein CD71, with an HA tag positioned C-terminally to the TMEM106B-CTD moiety (CD71-CTD-HA). This construct was introduced into the non-permissive BHK21 cells and resultant cells were subsequently inoculated with SARS-CoV-2-mNeon Green (mNG) S^E484D^. Immunoblotting and cell-surface HA staining confirmed expression and plasma-membrane localization of the CD71-TMEM106B-CTD chimera (**Fig. 6B, 6C**), although only ∼5% of the cells expressed CD71-TMEM106B-CTD due to limited antibiotic selection to minimize toxicity. Notwithstanding, at 24 hpi, a measurable number of BHK21-CD71-CTD-HA cells supported infection compared to WT BHK21 cells (**Fig. 6D**) as also evident in a 2-log increase in cell-associated viral RNA levels at 48 hpi (**Fig. 6E**). Infection of BHK21-CD71-CTD-HA cells was still mediated by an endosomal pathway, as inhibition of endosome acidification by ammonium chloride treatment reduced cell-associated viral RNA levels by ∼100-fold (**Fig. 6F**).

**Figure 6.**
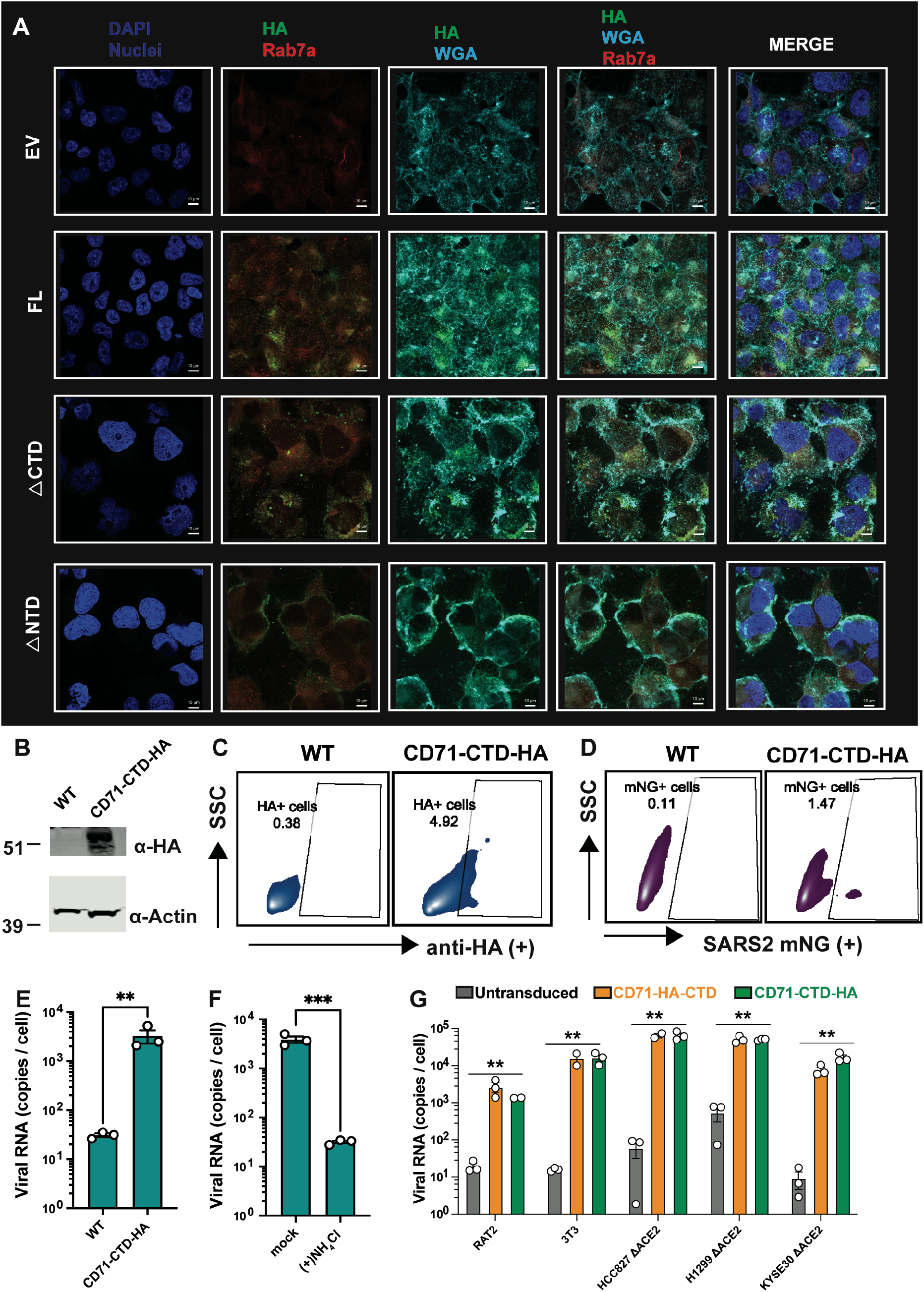
Targeting TMEM106B-CTD to the plasma membrane is sufficient to confer susceptibility to SARS-CoV-2 SE484D. **(A)** H522 TMEM106B KO cell derivatives stably transduced with an empty vector (EV), and HA-tagged full length TMEM106B (FL), TMEM106BΔCTD, TMEM106BΔNTD were subjected to immunofluorescence microscopy. Detection of TMEM106B variants (anti-HA, green), Rab7a (in red), wheat germ agglutinin (membrane, cyan), and cellular nuclei (DAPI, blue). Scale bar, 10 µm.μ **(B, C)** BHK-21 cells stably expressing CD71-TMEM106B-CTD-HA (CD71-CTD-HA) chimeric protein were subjected to western blotting (**B**) or surface staining and flow cytometry (**C**) using an anti-HA antibody. **(D, E)** BHK21-CD71-TMEM106B-CTD-HA cells were inoculated with SARS-CoV-2 mNG S^E484D^ at an MOI of 1. Infected cells were subjected to FACS to enumerate mNG positive cells at 48 h post-infection **(D)** or RT-qPCR analysis to analyze cell-associated SARS-CoV-2 RNA at 48 h post infection **(E).** (**F**) BHK21-CD71-CTD-HA cells were infected as in (D, E) but in the presence of 50 mM ammonium chloride. Cell-associated SARS-CoV-2 RNA levels were assessed by RT-qPCR at 48 h post infection. (**G**) Rat2, 3T3, and ACE2 KO HCC827, H1299 and KYSE30 cells were transduced with CD71-HA-CTD or CD71-CTD-HA constructs and infected with SARS-CoV-2 mNG S^E484D^ at an MOI of 1. Cell-associated SARS-CoV-2 RNA levels were assessed by RT-qPCR at 72 h post infection. Data in E-G show the average of two-three independent replicates with error bars displaying the SEM. \*\**P* < 0.01; \*\*\**P* < 0.001; by two-tailed unpaired *t*-test or one-way ANOVA with Dunnett’s correction for multiple comparisons.

Similar to BHK-21 hamster cells, expression of CD71-CTD-HA in non-permissive rat (RAT2), mouse (3T3) and human cells (HCC827, H1299 and KYSE30) engineered to lack ACE2 resulted in a significant increase in cell-associated viral RNA levels upon infection with SARS-CoV-2-mNG S^E484D^ (**Fig. 6G**). In addition, changing the position of the HA-tag to the N-terminus of CTD (CD71-HA-CTD) did not further enhance replication (**Fig. 6G**). Furthermore, addition of anti-HA antibodies to cultures of HCC827 and H1299 cells expressing CD71-HA-CTD or CD71-CTD-HA resulted in accumulation of the HA antibody at the plasma membrane as well as in late endosomes/lysosomes marked by LAMP1 (**Fig. 7A**). Similarly, addition of a TMEM106B-CTD-specific antibody to cultures of H522 cells expressing full-length TMEM106B or TMEM106B-ΔNTD also resulted in accumulation of the antibody in LAMP1 positive late endosomes/lysosomes (**Fig. 7B**). As TMEM106B is broadly expressed across various cell types, these findings indicate that the endogenous levels of TMEM106B present in non-permissive cells are not sufficient and that the presence of TMEM106B-CTD at the plasma membrane is required to allow SARS-CoV-2-mNG S^E484D^ replication.

**Figure 7.**
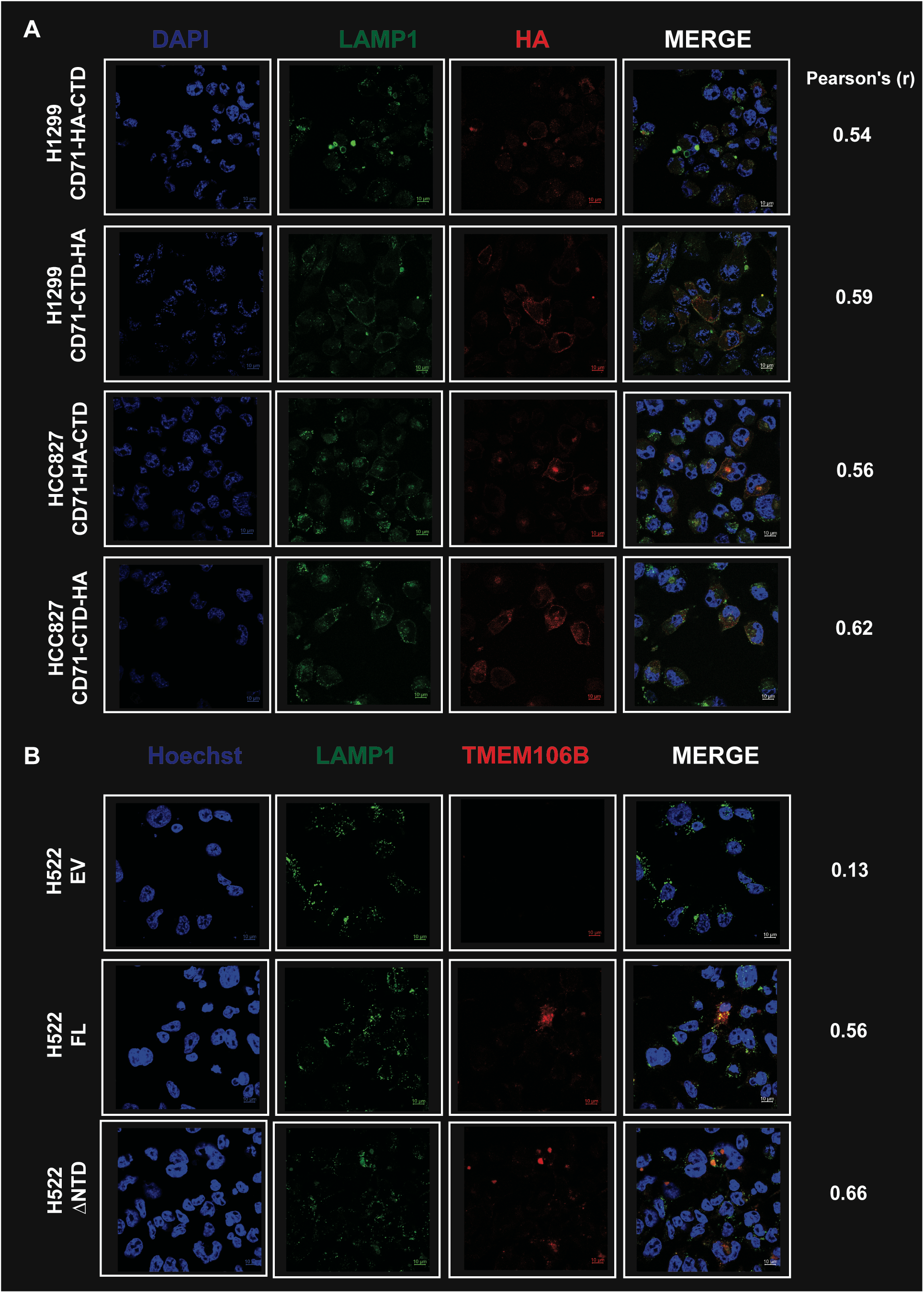
Membrane-targeted TMEM106B-CTD and full-length TMEM106B traffic from the plasma membrane to LAMP1-positive late endosomes/lysosomes. (A) H1299 and HCC827 cells stably expressing CD71-HA-TMEM106B-CTD (CD71-HA-CTD) or CD71-TMEM106B-CTD-HA (CD71-CTD-HA) were incubated with an Alexa Fluor 594-conjugated anti-HA antibody (red) for 2 h at 37°C, then washed, fixed, and permeabilized. Lysosomes were labeled with an anti-LAMP1 antibody (green) and nuclei were counterstained with DAPI (blue). (B) H522 cells stably expressing an empty vector (EV), TMEM106B-FL, or TMEM106B-ΔNTD were incubated with an anti-TMEM106B-CTD antibody (red) and processed as in (A), with lysosomes labeled by anti-LAMP1 (green) and nuclei counterstained with Hoechst (blue). Pearson’s correlation coefficients (r) between the LAMP1 and HA or TMEM106B channels, calculated in Fiji using the Coloc 2 plugin with automatic thresholding, are shown to the right of each row. Images were acquired on a Zeiss LSM 880 Airyscan confocal microscope in super-resolution mode. Scale bar, 10 µm. Pearson’s r values are the mean of 4 fields.

### Heparan sulfates are not required for TMEM106B-mediated entry at high expression levels of TMEM106B

TMEM106B is broadly expressed across all tissues but is enriched in the central nervous system (**Fig. 8A**). This raises the question as to why only a limited number of cell lines support TMEM106B-dependent infection. Given the dependency of TMEM106B-mediated entry on surface heparan sulfates (**Fig. 2E**), we first investigated whether cells expressing membrane-targeted TMEM106B also required heparan sulfate expression on their surface. Notably, heparinase-treated H522 cells expressing TMEM106B-ΔNTD supported SARS-CoV-2 S^E484D^ replication at similar levels as untreated cells, whereas WT H522 cells remained sensitive to heparan sulfate depletion (**Fig. 8B**). In agreement, ACE2 KO H1299, HCC827 and KYSE30 cells expressing membrane targeted TMEM106B were also largely insensitive to heparan sulfate depletion though we noted a statistically significant decrease in cell-associated viral RNA levels in KYSE30-ΔACE2-CD71-HA-CTD cells (**Fig. 8B**). As an alternative approach to heparinase treatment, we determined whether TMEM106B expression could restore replication in non-permissive PgsA-745 hamster cells that lack a functional xylosyltransferase enzyme, hence are deficient in glycosaminoglycan synthesis and lack surface heparan sulfate glycosaminoglycans. Expression of TMEM106B, TMEM106B-ΔNTD and membrane-targeted TMEM106B-CTD enhanced SARS-CoV-2 S^E484D^ replication in PgsA-745 cells, whereas TMEM106B-ΔCTD did not (**Fig. 8C, 8D**). Together, these data show that while surface heparan sulfates facilitate TMEM106B-mediated entry at endogenous expression levels, they are not required when TMEM106B is more abundantly present at the plasma membrane.

**Figure 8.**
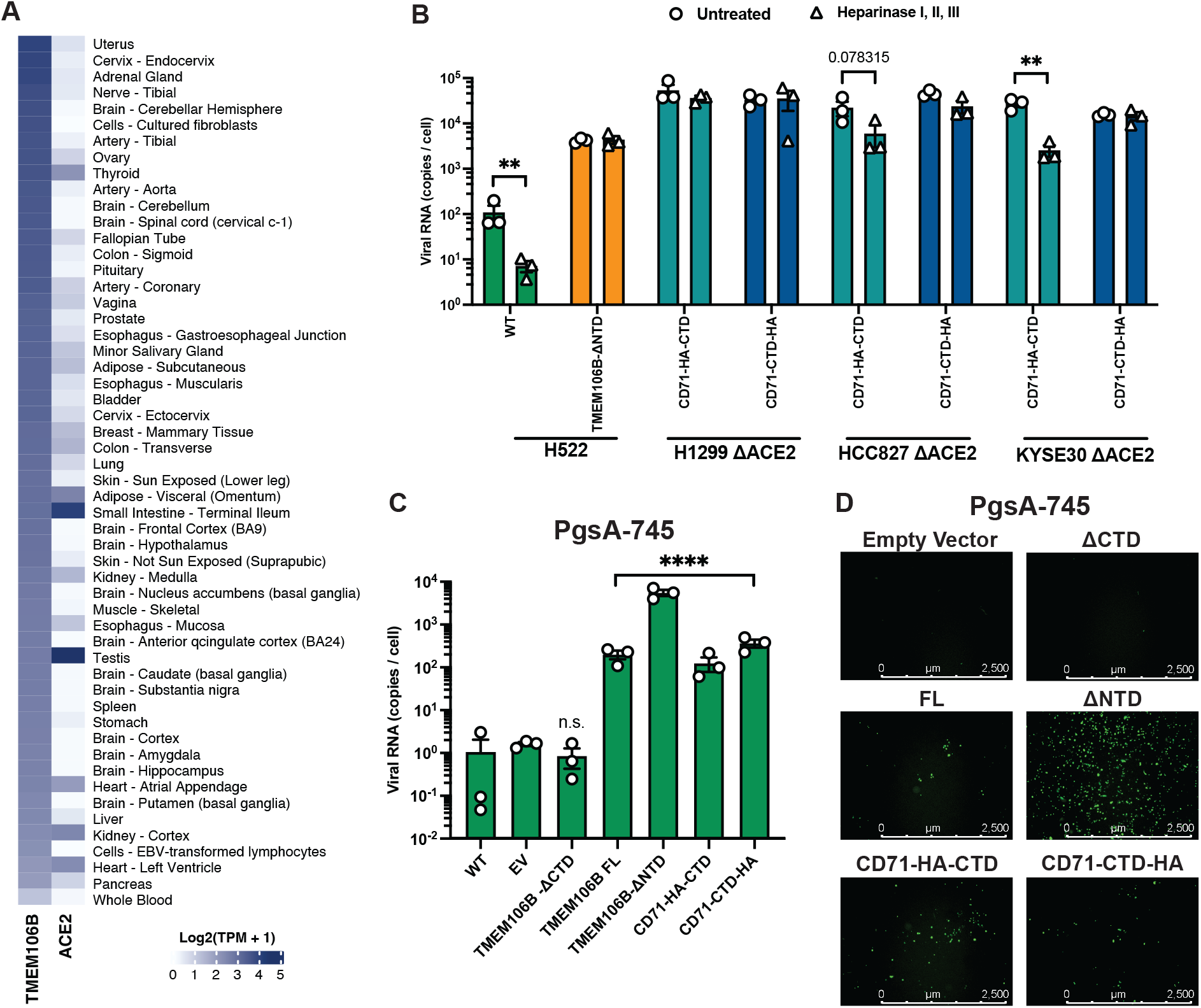
Heparan sulfates are dispensable for TMEM106B-mediated entry when TMEM106B-CTD is abundantly present at the plasma membrane. **(A)** Median bulk tissue expression of TMEM106B and ACE2 across human tissues, obtained from the Adult Genotype Tissue Expression (GTEx) Portal. **(B)** H522 WT and H522-TMEM106B-ΔNTD cells, and ACE2 KO H1299, HCC827, and KYSE30 cells expressing CD71-HA-TMEM106B-CTD (CD71-HA-CTD) or CD71-TMEM106B-CTD-HA (CD71-CTD-HA), were left untreated (circles) or treated with a combination of heparinase I, II, and III (triangles) for 90 min prior to inoculation with SARS-CoV-2-mNG S^E484D^ at an MOI of 1. Cell-associated viral RNA was quantified by RT-qPCR at 72 hpi. **(C)** PgsA-745 cells stably transduced with an empty vector (EV), TMEM106B-ΔCTD, TMEM106B-FL, TMEM106B-ΔNTD, CD71-HA-CTD, or CD71-CTD-HA were inoculated with SARS-CoV-2-mNG SE484D at an MOI of 1 and cell-associated viral RNA was quantified by RT-qPCR at 72 hpi. **(D)** Representative fluorescence microscopy images of mNG-positive PgsA-745 cells from the experiments shown in (C). Data in B and C show the mean of three independent replicates; error bars show the SEM. ns, not significant; **P < 0.01; ****P < 0.0001 by two-tailed unpaired t test or one-way ANOVA with Dunnett’s correction for multiple comparisons.

### Serial passaging selects for the S-V951A substitution which confers enhanced fusogenicity to S^E484D^ and allows enhanced ACE2-independent infection in H522 and H661 cells

Despite the strict requirement for the S-E484D substitution for ACE2-independent entry in H522 and H661 cells, we noted that infection rates were relatively low within the context of the VSV-SARS-CoV-2 S^E484D^ chimeric virus (**Fig. 9A**). Together with the observed low affinity of S^E484D^ for TMEM106B (**Fig. 5E**, see also (10)), this prompted us to test whether additional S substitutions could enhance utilization of the TMEM106B-dependent entry pathway. To this end, we serially passaged VSV-GFP-SARS-CoV-2-S^E484D^ in H522 cells. Viral titers progressively increased in culture supernatants, and later passages infected cells at substantially lower input volumes (**Fig. 9B**). Importantly, passaged viruses retained TMEM106B dependence (**Fig. 9B**) and showed similarly enhanced infectivity in H661 cells (**Fig. 9C**).

**Figure 9.**
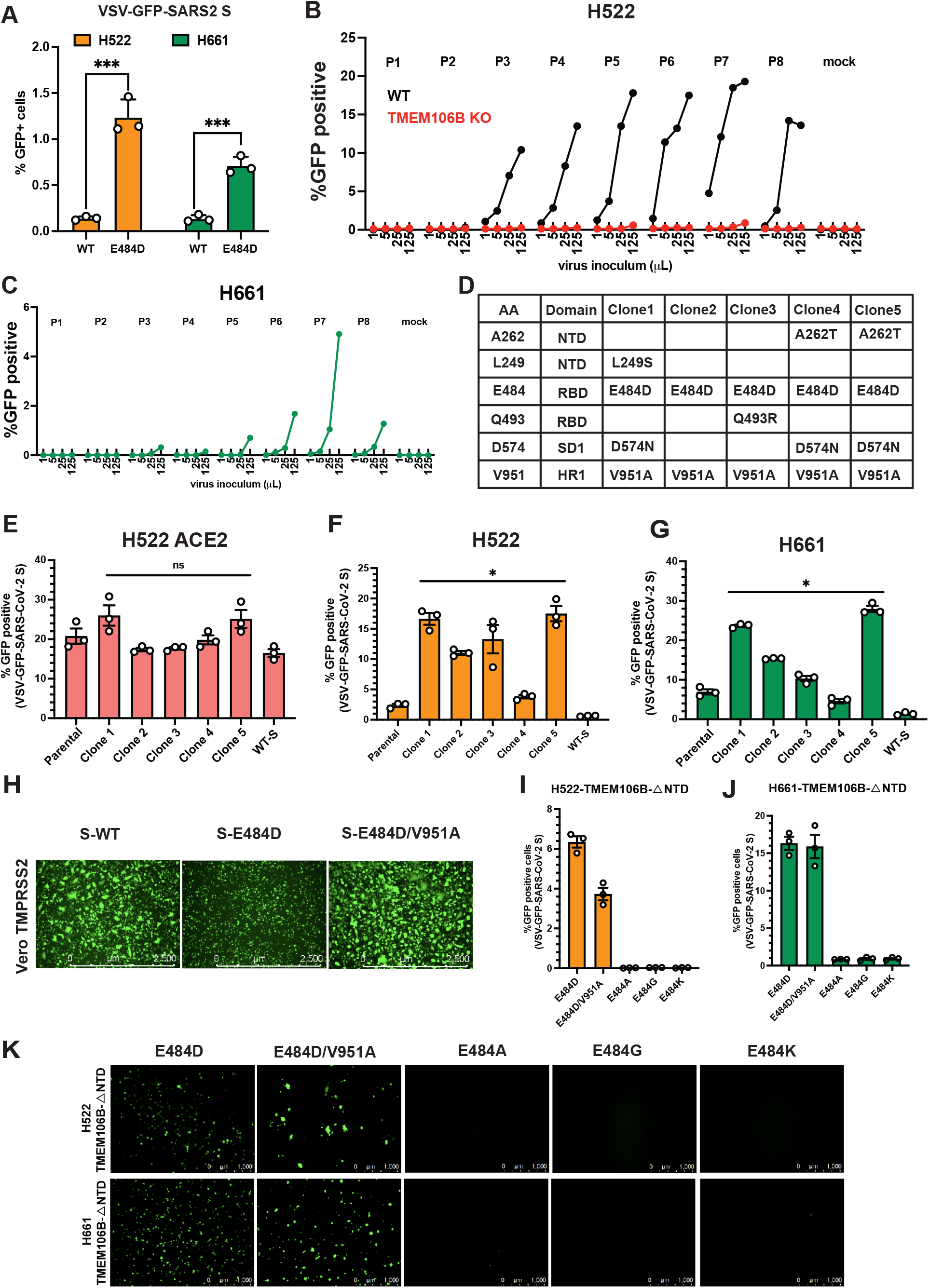
Serial passaging selects for an S-V951A substitution that enhances fusogenicity and ACE2-independent infection in H522 and H661 cells. (A) H522 and H661 WT cells were inoculated at an MOI of 1 with VSV-GFP-SARS-CoV-2 bearing WT S or the S^E484D^ substitution. GFP-positive cells were analyzed by flow cytometry at 48 hpi. (B) VSV-GFP-SARS-CoV-2 S^E484D^ was serially passaged in H522 cells for eight rounds. Virus from each passage (P1–P8) was titered on H522 WT (black) and H522-TMEM106B KO (red) cells using 1, 5, 25, or 125 µL of inoculum. The percentage of GFP-positive cells at each inoculum volume is shown. (C) Virus from each passage in (B) was titered on H661 cells as in (B). (D) Five virus clones were plaque-purified from passage 7 and sequenced. The table lists the observed amino acid substitutions and their location within S. (E–G) H522-ACE2 (E), H522 (F), and H661 (G) cells were inoculated at an MOI of 0.1 with parental VSV-GFP-SARS-CoV-2 S^E484D^, the five plaque-purified clones from (D), or VSV-GFP-SARS-CoV-2 bearing WT S. H522-ACE2 cells were fixed at 16 hpi and H522 and H661 cells at 48 hpi for enumeration of GFP-positive cells by flow cytometry. (H) Vero-TMPRSS2 cells were infected with VSV-GFP-SARS-CoV-2 bearing S-WT, S-E484D, or S-E484D/V951A at an MOI of 0.1, and syncytia formation was assessed by fluorescence microscopy at 24 hpi. (I, J) H522-TMEM106B-ΔNTD (I) and H661-TMEM106B-ΔNTD (J) cells were inoculated with VSV-GFP-SARS-CoV-2 bearing S-E484D, S-E484D/V951A, S-E484A, S-E484G, or S-E484K at an MOI of 0.1. GFP-positive cells were analyzed by flow cytometry at 48 hpi. (K) Representative fluorescence microscopy images of the cells in (I) and (J). Scale bar, 2,500 µm.μ Data in A and E–J show the mean of three independent biological replicates; error bars show the SEM. In E–G, all clones were compared to the parental virus. (ns, not significant; *P < 0.05; ***P < 0.001 by two-tailed unpaired t test or one-way ANOVA with Dunnett’s correction for multiple comparisons.)

Plaque purification and Sanger sequencing of five passage-7 isolates showed that all clones retained the E484D substitution while acquiring additional mutations across *S*, including changes in the NTD, RBD, SD1, and HR1 (**Fig. 9D**). We next assessed which substitutions contributed to enhanced infectivity. Passaged viruses infected H522-ACE2 cells with similar efficiency (**Fig. 9E**), indicating that these mutations did not impair ACE2-mediated entry or overall viral fitness. Except for Clone 4, which carried the same set of mutations as Clone 5, all clones displayed comparable increases in infectivity in both H522 WT and H661 WT cells (**Fig. 9F, 9G**). Notably, clone 2 —which harbored only one additional mutation (V951A)—also exhibited a similar level of enhanced replication (**Fig. 9F, 9G**), suggesting that the HR1 substitution V951A, present in all passaged clones, likely contributes to the increased virion infectivity.

V951 residue is located within the heptad repeat 1 (HR1) region which is essential for fusion pore formation (19). To examine whether the V951A substitution confers enhanced overall fusogenicity to S-E484D independent of TMEM106B, we examined syncytia formation in Vero-TMPRSS2 cells. Despite VSV-GFP-SARS-CoV-2 S^E484D^ showing no obvious replication defect in ACE2-expressing cells (**Fig. 5E**, see also (9)), we observed markedly less syncytia formation (**Fig. 9H**). In contrast, VSV-GFP-SARS-CoV-2 S^E484D/V951A^ double mutant formed syncytia at levels matching that observed with the WT virus (**Fig. 9H**). Of note, while the S-E484D/V951A double mutant did not increase the number of infected H522 and H661 cells ectopically expressing TMEM106B-ΔNTD (**Fig. 9I, 9J**), likely due to its high surface expression, we saw a similar tendency to form larger syncytia in these cells (**Fig. 9K**). None of the other VOC E484 mutations mediated infection of H522-TMEM106B-ΔNTD and H661-TMEM106B-ΔNTD cells (**Fig. 9I-K**). Collectively, these data reinforce the absolute necessity of the S-E484D substitution for TMEM106B-dependent infection and demonstrate that the S-V951A substitution enhances the overall fusogenicity of S independent of TMEM106B usage.

### E484D is mutationally accessible but rare in nature

To place E484D in a natural context, we analyzed E484 substitutions across publicly available data sets. Of the six amino acids reachable from the parental GAA codon in one nucleotide step, E484D is the only one reachable by two independent routes (23014 A>C or A>T), yet it is the rarest substitution found only in 229 out of 9,438,404 GenBank genomes (0.0024%, **Fig. 10A**) from LAPIS/CoV-Spectrum (20). In contrast, E484A (Omicron) is found in 45.6% of genomes, and E484K in 3.9% (**Fig 10A**). Substitutions that are as accessible as E484D but far rarer in nature point to selection, rather than mutational supply, as the limit on emergence of E484D. Amongst the 406 SRA data sets in which E484D was found, the median allele fraction was 0.98, and 330 runs had E484D above 50% (**Fig 10B**). Because only runs with E484D detected are included, these numbers describe the frequency of E484D within a sample where it was detected, not how often E484D occurs in circulating virus. The MA1 mouse-adaptation lineage, in which E484D reached 96% of reads in mouse lung and 92% in cell supernatant, is plotted for comparison from Yan et al. (21).

**Figure 10.**
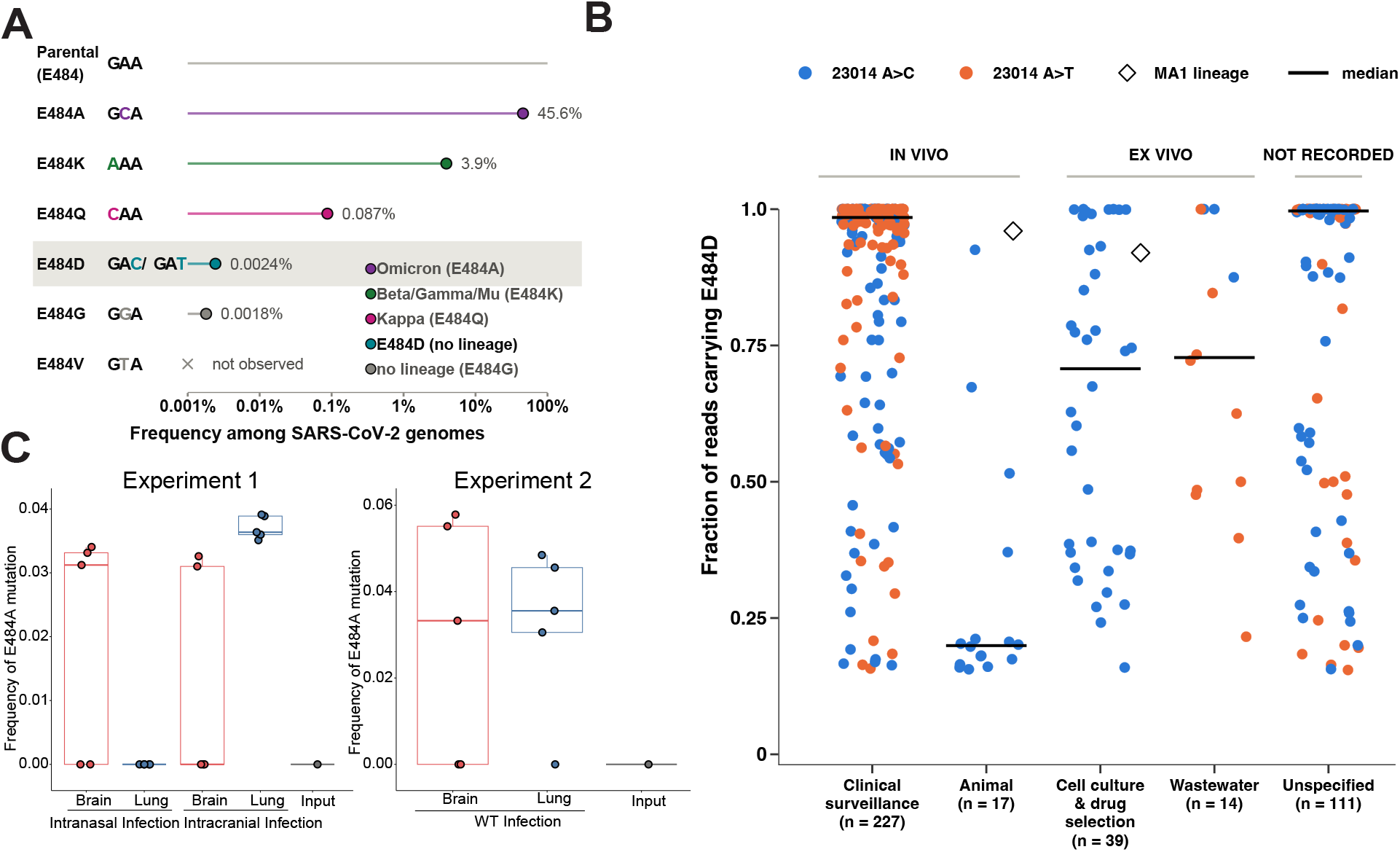
E484D is mutationally accessible but rare in nature whereas E484A arises *in vivo*. **(A)** Substitutions reachable from the parental GAA codon in one nucleotide step, with each changed base highlighted in the color of its lineage. Frequencies are the fraction of 9,438,404 GenBank genomes carrying each change, on a log scale. Colors mark lineage: Omicron (E484A), Beta/Gamma/Mu (E484K) and Kappa (E484Q). E484D and E484G have no lineage assignment, and E484V was not observed. **(B)** Fraction of reads carrying S-E484D in each SRA run (n = 406), grouped by sample context. In vivo: clinical surveillance, including one patient monoclonal-antibody treatment run; and animal, combining wild deer and experimental animals, with the MA1 mouse-lung point. Ex vivo: cell culture, including drug-selection passages (molnupiravir, remdesivir) and the MA1 cell-supernatant point; and wastewater. Unspecified sample context is shown separately. Each point is one run; horizontal bars show medians; diamonds mark the MA1 mouse-adaptation lineage, plotted from a different study. Blue and orange mark the 23014 A>C and A>T routes. **(C)** Frequency of the spike E484A substitution (A23013C) in viral RNA from the brain (red) and lung (blue) of mice inoculated intranasally or intracranially (Experiment 1, left) or only infected intranasally (Experiment 2, right). Input refers to sequenced viral inoculum of the reference WA strain of SARS-CoV-2 virus. Each symbol represents one animal; boxes show the median and interquartile range. Only positions with ≥100-fold coverage were analyzed; samples without detectable minority mutants are plotted at 0.

Because E484 is a recurrent site of antigenic variation in circulating SARS-CoV-2 lineages, we asked whether substitutions at this residue arose during replication in different tissue compartments, as we had previously reported for the furin cleavage site in a murine model (22). To this end, we analyzed published data sets from the CNS and lung, after either intranasal or intracranial infection. The E484A substitution was undetectable in the sequenced virus inoculum but was recovered as a minority variant at frequencies of approximately 3–6% of reads in a subset of animals in both experiments, indicating de novo emergence during *in vivo* replication rather than selection of a variant already present in the stock (**Fig. 10C**). The E484A mutation was recovered from brain and lung after both intranasal and intracranial inoculation and from both compartments in wild-type-infected animals, with overlapping frequency ranges between tissues (**Fig. 10C**). However, both experiments consistently showed higher diversity levels of this position in the brain. Notably, intracranially infected animals showed the highest levels of diversity (Fig. 10C), possibly indicating adaptive effects of this position during viral spread to new anatomical compartments.

## DISCUSSION

In this study, we define TMEM106B as an essential host determinant of ACE2-independent SARS-CoV-2 entry for the SARS-CoV-2 S^E484D^ variant. Using two independent lung-derived epithelial cell line models that support ACE2-independent infection, we demonstrate that the luminal TMEM106B-CTD mediates direct SARS-CoV-2 S^E484D^ binding and entry. Our findings extend recent reports that TMEM106B can support SARS-CoV-2 entry in the absence of ACE2 expression (12, 23, 24), provide mechanistic insight into how it functions in concert with surface heparan sulfates, and establish its functional requirement in cell types that use this alternative mechanism. More specifically, our findings indicate that at endogenous expression levels in H522 and H661 cells, likely a small fraction of TMEM106B localizes to the plasma membrane, which necessitates the additional use of heparan sulfates to capture SARS-CoV-2 S^E484D^ to mediate entry (**Fig. 11**). In contrast, if TMEM106B is more abundantly present at the plasma membrane, TMEM106B-CTD is sufficient to mediate virus binding and entry (**Fig. 11**).

**Figure 11.**
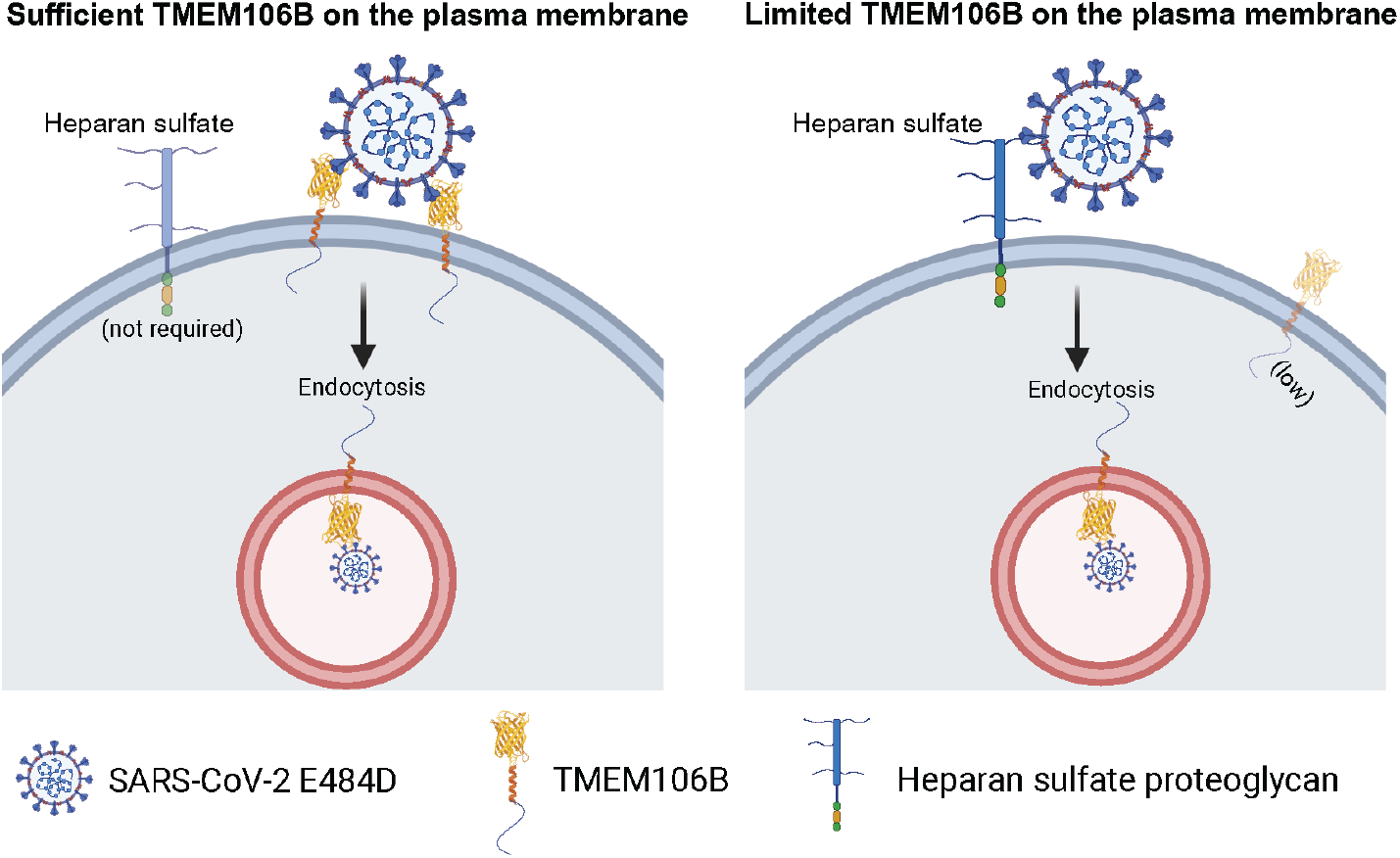
Model for TMEM106B-dependent, ACE2-independent entry of SARS-CoV-2 S^E484D^. When TMEM106B is abundant at the plasma membrane (left), SARS-CoV-2 S^E484D^ virions engage TMEM106B-CTD directly without the need for cell-surface heparan sulfate proteoglycans and are internalized by endocytosis. When limited amounts of TMEM106B are present at the plasma membrane, heparan sulfate proteoglycans are required to concentrate virions at the cell surface and promote engagement of TMEM106B. In both cases, fusion and genome release occur in an acidified endosomal compartment. Model was created with BioRender.com.

In contrast to prior reports suggesting that TMEM106B also enhances replication of a WT reference strain (10), we found a strict dependence on the S^E484D^ substitution for utilization of the TMEM106B-dependent entry pathway. This was most evident at the level of virus binding assays, whereby TMEM106B-ΔNTD markedly enhanced binding of SARS-CoV-2 S^E484D^ but not WT virus. Other spike substitutions at the E484 residue present in VOCs (e.g. E484A, E484G, E484K) also failed to support infection in the ACE2-independent cell models. Future studies will be required to determine whether additional S substitutions, alone or in combination, enable utilization of the TMEM106B-dependent entry pathway.

Whether the S^E484D^ substitution has relevance for disease pathogenesis is currently unclear. Notably, S^E484D^ substitution can arise as easily as other substitutions that became dominant in VOCs via a single mutation, suggesting that the virus must sample this substitution *in vivo*. Despite the lack of a replication defect in ACE2-expressing cells, our finding that VSV-GFP-SARS-CoV-2 S^E484D^ causes less syncytia in ACE2-and TMEM106B-expressing cells may suggest that it spreads largely cell-free. Cell-to-cell transmission may provide a fitness advantage within the mucous-rich airways that limit virus replication (25–27) and may explain why the seemingly subtle E484D substitution did not become fixed in circulating VOCs.

Nonetheless, the E484 residue has frequently mutated in VOCs, including E484K, E484Q and E484A (Omicron) as well as other less frequent substitutions (G, R, S, T), some of which are associated with immune escape (28). Our findings suggest that if the S^E484D^ becomes more prevalent, perhaps with an additional compensatory substitution such as V951A that allows more efficient cell-to-cell spread, it could alter the cell and tissue tropism of SARS-CoV-2 in humans. Of note, S^E484D^ substitution is frequently present across unique evolutionarily advanced SARS-CoV-2 cryptic lineages sampled from wastewater (29). These cryptic lineages are thought to represent individuals with very long persistent infections (possibly long COVID) raising the possibility that S^E484D^ arising within an individual might alter SARS-CoV-2 disease pathogenesis. As sequencing of SARS-CoV-2 S is typically done from upper airway samples, whether variation in S is associated with complex SARS-CoV-2 pathogenesis in other organs and long-term infection is unknown. In fact, a recent study in mice demonstrated higher levels of viral divergence in the CNS that is surprisingly not captured in sequences obtained from the lung (22). In addition, S^E484D^ confers resistance to mAb therapy in humans and naturally evolves in a long-term mouse xenograft infection model (28), further suggesting the potential *in vivo* relevance of this specific mutation.

Our studies support the direct involvement of TMEM106B in SARS-CoV-2 binding and suggest that at least a fraction of it must be present at the plasma membrane to mediate entry (**Fig. 11**). Although clustering of heparan sulfates may be sufficient for virus docking (30), complete loss of virion binding in TMEM106B KO cells argues against the model that heparan sulfates alone can mediate stable virus attachment. Rather, it is more likely that heparan sulfates enhance virus binding mediated by a small amount of TMEM106B present at the plasma membrane, in a manner similar to previous reports of heparan sulfates enhancing ACE2-dependent entry (31). While the enhanced ability of TMEM106B-ΔNTD to support virion binding correlated with its more prominent localization at the plasma membrane and targeting TMEM106B-CTD to the plasma membrane restored virus replication even in the absence of heparan sulfates (**Fig. 8B-D**), it is notable that viral entry in the latter case was still dependent on endosome acidification. TMEM106B primarily localizes to late endosomes and lysosomes (32, 33), and several intraluminal cues—acidic pH, proper glycosylation, membrane lipid composition, or endosome-restricted cofactors including cathepsins—may be required to support productive S^E484D^ engagement, reinforcing the idea that TMEM106B mediates a late-entry event (34).

Although TMEM106B is broadly expressed across tissues and cell types, it remains unclear why only H522 and H661 cells support TMEM106B-dependent entry. One possibility is that these cells have a greater proportion of TMEM106B at the plasma membrane, potentially due to differences in TMEM106B trafficking and expression levels, which can be modulated by N-glycosylation states, H⁺-ATPase activity, and TMEM106B mutations (22, 32). In addition, the types and levels of cell-surface heparan sulfates, which may act in concert with TMEM106B, could contribute to this cell-type specificity (35). We therefore propose that differences in cell-surface TMEM106B abundance and the composition and levels of heparan sulfate glycosaminoglycans underlie the restricted ability of certain cell types to support TMEM106B-mediated entry.

### Limitations of the study

A major limitation of our study is our inability to evaluate whether the S-E484D and TMEM106B-dependent entry pathway is physiologically relevant, which would require extensive characterization of SARS-CoV-2 variants within extra-respiratory tissues in humans. Of note, while the S-E484D substitution enhances neuroinvasiveness in mouse organoids, the presence of TMEM106B alone is not sufficient for SARS-CoV-2 replication in vivo (12). We cannot rule out whether knockout of TMEM106B, which has key functions in the lysosome, indirectly affects SARS-CoV-2 replication. However, complete inhibition of virus binding in TMEM106B KO cells, sufficiency of TMEM106B-CTD to mediate virus replication in non-permissive cells and inhibition of SARS-CoV-2 S^E484D^ replication by Fc-TMEM106B decoy together demonstrates the direct involvement of TMEM106B in virus binding and entry. At a molecular level, whether TMEM106B is sufficient for viral entry and TMEM106B-mediated entry depends on other co-factors (e.g. C-type lectin receptors (11)) and S-activating proteases such as TMPRSS2 and cathepsins in our cell models will also need to be addressed in the future. Several host attachment factors have been reported to facilitate ACE2-dependent or ACE2-independent SARS-CoV-2 entry (36), including Neuropilin 1 (37, 38), heparan sulfates (39), C-type lectin receptors (40–42), phosphatidylserine receptors TIM-1, TIM-4 and AXL (43, 44), LDLRAD3, TMEM30A, and CLEC4G (45). Whether these factors function as physiologically relevant entry mediators at endogenous expression levels remains to be determined.

In summary, our work establishes TMEM106B as a key host entry factor enabling ACE2-independent infection by the SARS-CoV-2 S^E484D^ variant in H522 and H661 cells. The essential role of the TMEM106B CTD and the viral adaptations in S that optimize the utilization of TMEM106B provide insight into the mechanism of ACE2-independent SARS-CoV-2 entry in airway cell line models. Notably, TMEM106B is highly expressed in neurons, oligodendrocytes, and glial cells (32, 33) and altered TMEM106B expression and processing has been linked to several neurodegenerative diseases (13). This raises the possible hypothesis that TMEM106B-mediated entry could function as a SARS-CoV-2 entry factor in tissues such as the CNS, where ACE2 expression is limited (46, 47). Whether TMEM106B contributes to SARS-CoV-2 neuropathology or long-COVID remains an area for future investigation (48). Overall, our findings highlight that emerging mutations in viral S can allow the utilization of alternative entry pathways and broaden its cell and tissue tropism, which may help explain disease manifestations beyond the respiratory tract. Broadly, identification of alternative entry factors will inform our understanding of disease manifestations of emerging SARS-CoV-2 variants, and possibly related coronaviruses.

## MATERIALS AND METHODS

### Plasmids

pLentiCRISPRv2 (Addgene ID #98290) derived vector targeting *ACE2* has been described before (9). pLentiCRISPRv2 targeting *TMEM106B* was similarly generated. In brief, the pLentiCRISPRv2 plasmid was digested with BsmBI for 2 h at 55°C. Designed oligos (5’-TTTCAGAAAGCTGTATGTGA-3’ and 5’-AAAGTCTTTCGACATACACT-3’) were phosphorylated with T4 PNK at 37°C for 30 min and annealed. The resulting DNA was diluted 1:200 with H_2_O, ligated to the digested vector, and transformed into DH10B cells. The presence of the designed sgRNA oligos was verified by Sanger sequencing.

Gene fragments encoding full-length *Homo sapiens* transmembrane protein 106B (TMEM106B), TMEM106B-ΔNTD (Δ6-94aa) and TMEM106B-ΔCTD (Δ123-274) were synthesized as gBlocks by Integrated DNA Technologies (IDT) and amplified using the primers listed in Table 1. Briefly, the TMEM106B-ΔCTD construct terminates five amino acids downstream of the transmembrane (TM) alpha-helix, while the TMEM106B-ΔNTD construct retains the TM alpha-helix with the N-terminal domain deleted (residues 6–94 upstream of the domain). All constructs also contained an N-terminal HA epitope tag appended to the TMEM106B sequence via a Gly-Ser (GGS) linker. pCAGGS vector was linearized with SacI and NheI, and the amplified fragments were inserted using a homology-based cloning system (Takara Bio) following the manufacturer’s protocol. The plasmid sequences were confirmed by deep sequencing (Plasmidsaurus). The resultant pCAGGS-HA-TMEM106B-FL, pCAGGS-HA-TMEM106B-ΔCTD, pCAGGS-HA-TMEM106B-ΔNTD plasmids were subsequently used as template in PCR reactions using the primers listed in Table 1. These products were then inserted into the pLV-EF1a-IRES-NeoR/KanR vector (Addgene #85139) that has been digested with BsmBI and BamHI using quick ligation (NEB). The sequence of the resulting construct was confirmed by Sanger sequencing and propagated in DH10B *E. coli* cells (Life Technologies).

**Table 1:**
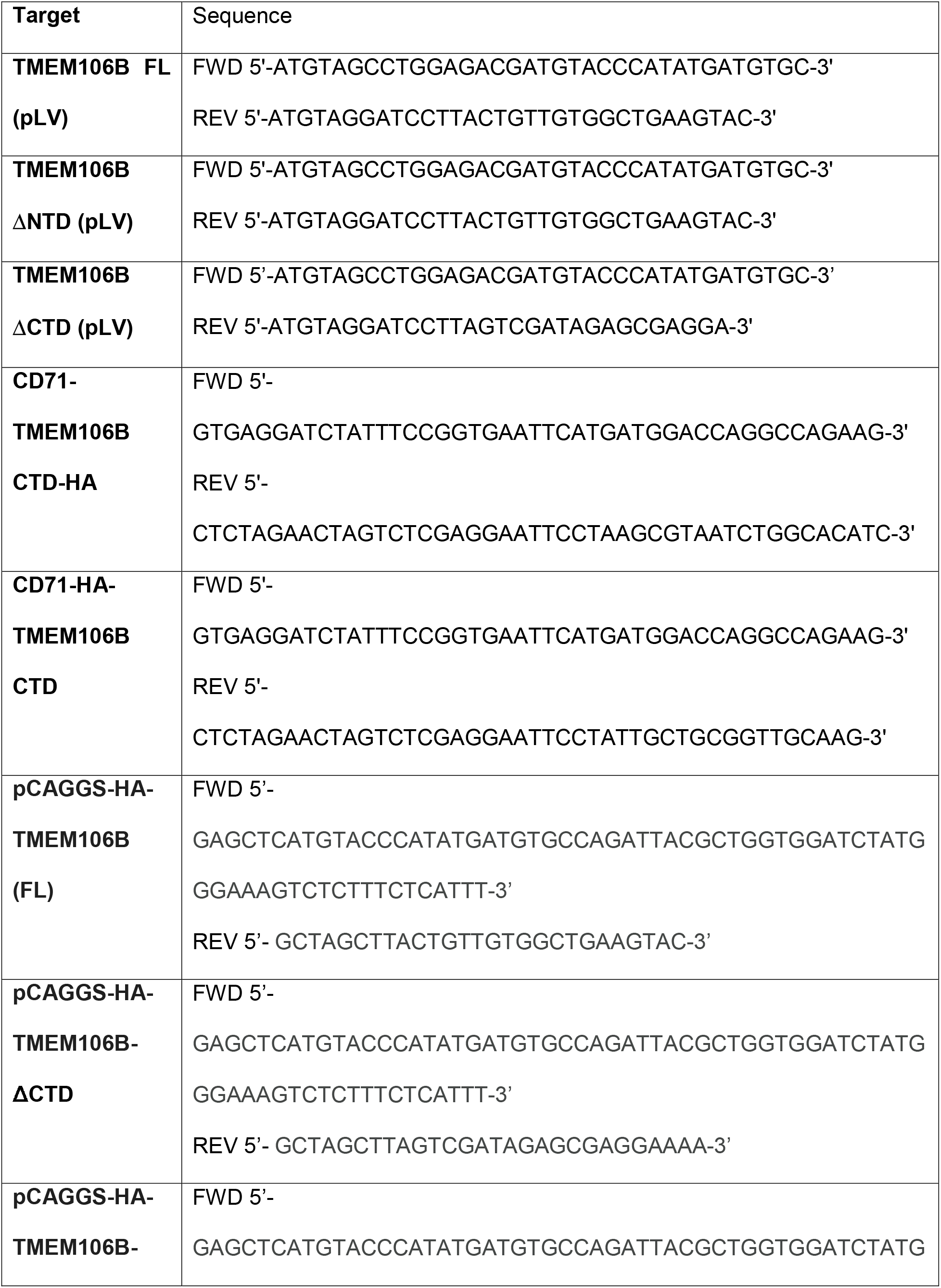
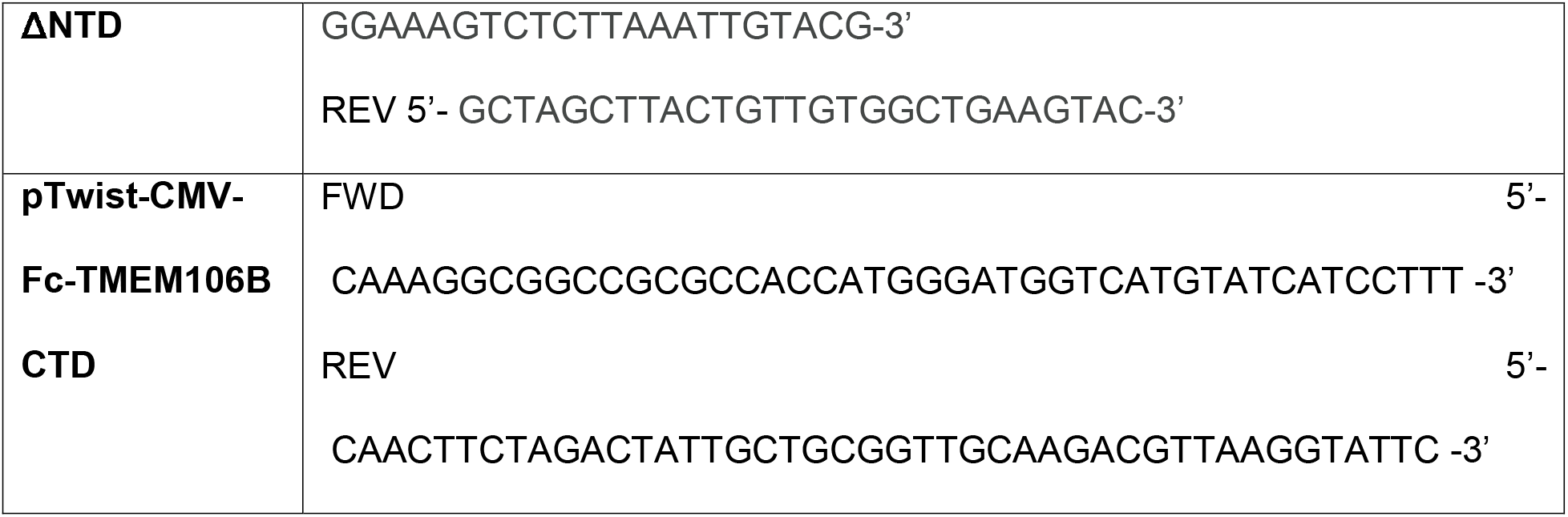
List of primers used in cloning of TMEM106B and derivatives.

Gene fragments encoding the CD71-TMEM106B-CTD-HA and CD71-HA-TMEM106B-CTD chimeras were ordered from Integrated DNA Technologies (IDT) as gBlocks. In both constructs, the N-terminal cytoplasmic and transmembrane domains of human transferrin receptor 1 (CD71/TFRC, residues 1-121) were joined through a Gly-Gly-Ser-Gly-Gly-Ser (GGSGGS) linker to the C-terminal luminal domain of TMEM106B (amino acids 118-274), replacing the native CD71 ectodomain so that the TMEM106B-CTD is displayed on the extracellular face of the plasma membrane. The two constructs differ only in the position of the HA epitope tag (YPYDVPDYA), which was placed either immediately downstream of the TMEM106B-CTD (CD71-TMEM106B-CTD-HA) or between the GGSGGS linker and the TMEM106B-CTD (CD71-HA-TMEM106B-CTD); both encode a 293-amino-acid protein. PCR amplified (see Table 1) and restriction enzyme digested fragments were similarly inserted into the pLV-EF1a-IRES-Blast (Addgene #85133) backbone. Insert identity and junctions were confirmed by Sanger sequencing.

The pTwist-hIgG1-TMEM106B-CTD was designed as a soluble receptor decoy to inhibit viral entry similar to approaches as described before (49, 50). In brief, an N-terminal domain with the mouse IgK heavy chain signal peptide (MGWSCIILFLVATATGVHS) followed by a GGS linker was included as a secretion-competent scaffold to promote efficient folding and extracellular expression of the fusion protein. This domain is followed by the Fc region of human IgG1, which mediates disulfide-linked dimerization, increases protein stability, and enables purification via Protein A/G affinity chromatography. A small, flexible peptide linker (GGS) separates the Fc domain from the C-terminal luminal domain of TMEM106B (amino acids 118-275), allowing independent folding and accessibility of the receptor-binding interface. The TMEM106B luminal domain serves as the viral binding module, enabling the secreted Fc fusion protein to function as a soluble decoy that sequesters virus particles and competitively blocks TMEM106B-mediated viral entry.

### Cell lines

The NCI-H522 (ATCC-CRL-5810) and NCI-H661 (ATCC-HTB-183) lung adenocarcinoma cells, the NCI-H1299 (ATCC-CRL-5803) non-small cell lung carcinoma cells, KYSE30 cell line (kind gift from the John Hayes Lab (University of Tennessee Health Science Center)) and HCC827 (ATCC CRL-2868) (kind gift from Luke Chen (North Carolina Central University)) cell lines were cultured in RPMI-1640 (Corning) supplemented with 10% FBS (Sigma) and 1% penicillin–streptomycin (Corning). PgsA-745 (ATCC #CRL 2242) cells were cultured in DMEM/nutrient mixture F-12 Ham Medium, supplemented with 10% FBS (VWR). BHK-21 (ATCC-CCL-10) cells were grown in DMEM: F12 Medium supplemented with 10% FBS. Rat2 (ATCC-CRL-1764) and NIH/3T3 (ATCC-CRL-1658) cells were cultured in Dulbecco’s Modified Eagle’s Medium (DMEM, Sigma), supplemented with 10% FBS. Vero E6 cells (ATCC-CRL-1586) were cultured in Minimal Essential Medium (MEM, Sigma), supplemented with 10% FBS and 10 mM HEPES buffer.

### Bioinformatics analyses

For selection of H661 cells as an ACE2-independent cell line model, bulk RNA-seq expression profiles (TPM) and metadata for human cancer cell lines were obtained from the DepMap portal (51). Cell lines were filtered to those with low ACE2 expression (log₂(TPM+1) ≤ 0.165) and low interferon-γ pathway activity (normalized enrichment score < 1), with interferon-γ activity quantified by gene set enrichment analysis (GSEA) of the *reactome_interferon_gamma_signaling* gene set. To identify cell lines with cell-surface expression patterns similar to H522, expression profiles were restricted to genes annotated for localization to the cell surface or plasma membrane and embedded using UMAP. Candidate cell lines were evaluated based on proximity in surface-expression space to H522, and H661 was selected based on strong surfaceome similarity while meeting the combined ACE2 and IFN-γ filtering criteria.

For analysis of TMEM106B and ACE2 expression across tissues, median bulk tissue gene expression values (TPM) for *TMEM106B* and *ACE2* were obtained from the Adult Genotype Tissue Expression (GTEx) Portal (v10) on 04/29/2026. Values were log-transformed (log₂[TPM + 1]).

### Heparinase treatment

We removed cell surface heparan sulfates using a combination of heparinase I (5.8 µg/mL), II (11 µg/mL) and III (11.2 µg/mL) from two vendors Sigma Scientific (#H3917) and Bio-techne R&D systems (#7897-GH (heparinase I), 6336-GH (heparinase II), 6145-GH (heparinase III)). Cells were seeded in 24-well plates and treated with 200 µL of media containing heparinases (heparinase I, 5.8 µg/mL; heparinase II, 11 µg/mL; heparinase III, 11.2 µg/mL) for 90 min at 37°C. Virus inoculum (MOI of 0.1/cell) was added and incubated for 2 h at 37°C for adsorption. Virus inoculum and heparinases were removed, and wells washed with 1X PBS and replaced with RPMI-10%FBS. Cell associated viral RNA was quantitated as described below.

### CRISPR-Cas9 knockout cell lines

H661, KYSE30, HCC827, H1299 cells were transduced with a pLentiCRISPRv2 derived vector targeting ACE2. The sgRNA (GTACTGTAGATGGTGCTCAT) targets exon 3 of ACE2 and has been described before (9). After transduction, cells were selected in 2 μg/mL of puromycin. Clonal H661 ACE2 KO cells were isolated after limiting dilution in 96-well plates. Knockout status of isolated clones was confirmed by Sanger sequencing. H522 and H661 cells were transduced with a pLentiCRISPRv2 derived vector targeting *TMEM106B*. The sgRNA (TTTCAGAAAGCTGTATGTGA) targets exon 2 of *TMEM106B*. After transduction, H522 and H661 TMEM106B KO cells were selected in 0.5 µg/mL of puromycin and maintained as bulk populations. In some experiments, these cells were complemented with empty vector, full length (FL) TMEM106B, TMEM106B-ΔCTD and TMEM106B-ΔNTD as detailed below.

### Stable expression of TMEM106B and its derivatives, lentivirus production and transduction

Cell lines stably expressing TMEM106B and its derivatives (TMEM106B-FL, TMEM106B-ΔNTD, TMEM106B-ΔCTD, CD71-TMEM106B-CTD-HA and CD71-HA-TMEM106B-CTD) were generated by lentiviral transduction, as described before (9). Briefly, lentiviruses were produced in HEK293T cells by polyethyleneimine (Polysciences)-mediated transfection using a pLV-EF1a-IRES-NeoR/KanR (Addgene #85139)-based vector expressing TMEM106B derivatives (FL, ΔNTD, ΔCTD), or a pLV-EF1a-IRES-Blast (Addgene #85133)-based vector expressing the CD71-TMEM106B-CTD chimeras, psPAX2 as packaging plasmid (Addgene #12260) and VSV-G expressing envelope plasmid (Addgene #12259).

Cell culture supernatants containing lentiviruses expressing TMEM106B derivatives (FL, ΔNTD, ΔCTD) were collected and used to transduce H522-TMEM106B KO, H661-TMEM106B KO, PgsA-745. Cell culture supernatants containing lentiviruses expressing CD71-TMEM106B-CTD chimeras were collected and used to transduce PgsA-745, BHK-21, 3T3, RAT2, KYSE30, HCC827 and H1299 ACE2 KO cells. Two days post-transduction, cells expressing TMEM106B derivatives (FL, ΔNTD, ΔCTD) were selected for two weeks in G418 (geneticin, 100 µg/mL), whereas cells expressing the CD71-TMEM106B-CTD chimeras were subjected to limited selection in 2 µg/mL blasticidin to minimize toxicity, to generate bulk populations expressing the indicated transgenes.

Cell surface expression of CD71-TMEM106B-CTD-HA was confirmed by anti-HA-surface staining. In brief, cells were washed with 1xPBS and detached by treatment with 1X PBS supplemented with 5mM EDTA. Collected cells were sequentially incubated in 1% BSA for 1 h, anti-HA (Biolegend, clone 16B12, 1:500) for 2 h and secondary antibody solutions (Alexa Fluor 488 1:2000) for 1 h. Antibodies were prepared in 1% BSA, and all incubations took place at room temperature. Cells were subsequently fixed by 4% PFA for 10 min and analyzed by flow cytometry.

### Viruses and infections

SARS-CoV-2 (USA-WA1-2020) virus stocks were grown in Vero-TMPRSS2 cells and plaque assays were used to determine the virus titer as before (9, 15). Vero-TMPRSS2 cells were inoculated with 10-fold serial dilutions of virus stocks, incubated for 1 h at 37°C with intermittent rocking, followed by addition of 2% methylcellulose and 2X MEM containing 4% FBS. 3 days post infection cells were fixed using 4% paraformaldehyde (PFA) and stained with crystal violet solution. Plaques from three virus dilutions were counted to determine virus titers.

For SARS-CoV-2 and VSV–GFP–SARS-CoV-2-S infections, cells were seeded at 70-80% confluency one day prior to infection. Infections were performed by addition of virus inoculum in cell culture media supplemented with 2% FBS and intermittent rocking for 2 h. Virus inoculum was removed, cells washed twice with 1x phosphate-buffered saline (PBS) and plated in cell culture media containing 10% FBS. Samples were subsequently processed for RNA extraction (described below) or flow cytometry (FACS) analysis. For FACS analysis of SARS-CoV-2 mNG or VSV-GFP-SARS2 S infected cells, cells were detached by trypsinization and fixed with 4% PFA for 20 min at room temperature. FACS was performed using a BD LSR Fortessa flow cytometer and analyzed using the FlowJo software.

VSV–GFP–SARS-CoV-2-S chimeric viruses and derivatives were published previously. The parental virus stock bearing the S^E484D^ substitution was serially passaged in H522 WT cells for eight rounds. At each passage, viral supernatants were collected, and half of the harvested amount was inoculated onto fresh cells while the remaining half was archived. As virus replication peaked at Passage 7, this virus stock was used to isolate viral clones by plaque purification on Vero E6 cells. Viral RNA from individual plaques was extracted by Trizol, reverse transcribed, PCR amplified and subjected to Sanger sequencing as done before (9).

### Purification of Fc-TMEM106B CTD decoy and blocking experiments

To express Fc-TMEM106B-CTD, plasmid constructs were transfected into Expi293 cells using manufacturer instructions. Harvested cell culture supernatants containing Fc-TMEM106B was clarified by centrifugation (4,000 x g) and passed through a 0.22-µm filterμ. Fc-TMEM106B-CTD fusion protein was bound to Protein A Sepharose on a gravity column. The column was washed with 50 volumes of 1x TBS (20 mM Tris pH 8.0, 150 mM NaCl). Proteins were eluted with Pierce Gentle Ag/Ab Elution Buffer, pH 6.6 (Thermo Fisher) and desalted into 1x TBS (20 mM Tris pH 8.0, 150 mM NaCl) using a PD-10 column (Cytiva). Depending on the yield, protein was concentrated with an Amicon 10 kDa centrifugal filter (Millipore). Protein purity was assessed by reducing and nonreducing SDS-PAGE followed by staining with SimpliSafe Coomassie reagent (Thermo Fisher). Gels were imaged with an iBright 1500 instrument (Thermo Fisher).

For Fc-TMEM106B-CTD decoy blocking experiments, 5×10^5^ H522 and H661 cells were seeded on 24-well plates. The following day, SARS-CoV-2 SE484D (MOI of 2 i.u./cell) was incubated with dilutions of Fc-TMEM106B-CTD in serum-free DMEM at 37°C for 30 min°. The mixture was subsequently inoculated onto H522 and H661 cells and incubated at 37°C for 2 h°. Inoculum was removed and washed with 1XPBS. Cell-associated viral RNA was analyzed 72 hours post-infection by RT-qPCR assays.

### RNA extraction and RT-qPCR

Cell-associated RNA was extracted by Trizol (Thermo Fisher Scientific) following manufacturer’s instructions and subjected to RT-qPCR analysis for viral and cellular RNA as detailed before (52). For quantification of viral RNA, 10 μL of the extracted RNA was used in a TaqMan-based RT-qPCR assay using TaqMan™ RNA-to-CT™ 1-Step Kit (Applied Biosystems, #4392938), alongside RNA standards, targeting SARS-CoV-2 N gene. The primers and probe sequences are as described before (52). For ACE2 mRNA quantification, RT-PCR was performed on 1μg of RNA using the iScript gDNA Clear cDNA Synthesis Kit (Bio-Rad) and analyzed by qPCR using PowerUp SYBR Green Master Mix (Applied Biosystems) on a QuantStudio 5 machine. RNA levels were quantified using the ΔC_T_ method with *RPL13a* as the reference target as described before (9).

### Immunoblotting

Cells were grown to 70% confluence and lysed in 1X radioimmunoprecipitation assay buffer (10% glycerol, 50 mM Tris-HCl pH 7.4, 150 mM NaCl, 2 mM EDTA, 0.1% SDS, 1% NP40, 0.2% sodium deoxycholate) containing protease inhibitors (Sigma). Proteins were separated by SDS-PAGE, transferred to nitrocellulose membranes, incubated in blocking buffer (LI-COR), and subsequently with primary antibodies in blocking buffer supplemented with 0.1% Tween-20 overnight at 4°C. Washed membranes were incubated for 2 h at room temperature in secondary antibody solution (LI-COR IRDye 680, 800; 1:10,000 in blocking buffer (LI-COR) supplemented with 0.1% Tween-20), imaged on an Odyssey® CLx system, and analyzed using Image Studio Software. Antibodies were used at the following dilutions: β-actin (Santa Cruz, SC-8432, 1:1000; Sigma #A5316, 1:5000), anti-HA (Biolegend, clone 16B12, 1:2000), ACE2 (R&D Systems #AF933, 1:200).

### Immunofluorescence microscopy

Cells grown on coverslips (Neuvitro, #GG-12-1.5-Collagen) were washed with Hank’s balanced salt solution (HBSS, Life Technologies, #14175-079) and stained with 200 µL of 5 µg/mL of wheat germ agglutinin conjugate (WGA, Life Technologies, Alexa Fluor^TM^ 633, #W21404) for 10 min at 37°C. Cells were then washed twice with HBSS and fixed with 500 µL of 4% paraformaldehyde at 37°C for 15 min. Subsequently, coverslips were washed twice with 500 µL of 1X HBSS and permeabilized with 200 µL 0.1% saponin for 10 min at room temperature. After permeabilization, cells were incubated in 1% bovine serum albumin (BSA) and 10% FBS in PBS containing 0.1% Tween-20 (PBST) at room temperature for 1 h. Samples were then incubated in primary mouse monoclonal anti-HA (Biolegend, clone 16B12, 1:500) and rabbit polyclonal anti-Rab7a (Novus Biologicals #NBP1-87174, 1:500) antibodies at 4 °C overnight. After washing with 1X HBSS, samples were incubated in a goat anti-mouse secondary antibody conjugated to Alexa Fluor Plus 488 (Invitrogen, Cat# A-11029, 1:1000) and goat anti-rabbit secondary antibody conjugated to Alexa Fluor Plus 568 (Invitrogen, Cat# A-11036, 1:1000) at room temperature for 1 h. Nuclei were stained with DAPI diluted in 1X HBSS at room temperature for 10 min. Finally, coverslips were washed in 1X HBSS and then mounted on slides using Prolong Gold Antifade. For confocal microscopy, samples were analyzed with a Zeiss LSM880 laser scanning confocal microscope (Carl Zeiss Inc. Thornwood, NY). Plan-Apochromat 20X numerical aperture (NA) 0.8), Plan-Apochromat 40X (NA 1.4) DIC oil, and Plan-Apochromat 63X (NA 1.4) DIC oil objectives were used. ZEN black software (version 2.1 SP3) was used for multichannel image acquisition and for obtaining Z stacks with optimal interval and section thickness. The image analysis software Volocity with Visualization module (version 6.3) (PerkinElmer, Waltham, MA) was used for the three-dimensional rendering of Z slices acquired through the depth of the cell monolayers.

### Virus attachment assays and RNAscope (FISH)

H522 and H661 cells (and their derivatives) were plated on 1.5 mm collagen-treated coverslips (GG-12-1.5-Collagen, Neuvitro) placed in 24-well plates one day prior to infection. For virus binding assays, cells were inoculated with SARS-CoV-2 S^E484D^ or SARS-CoV-2 WT-S at 16°C and fixed with 4% PFA 1 h post inoculation. Subsequently, cells were dehydrated with ethanol and stored at –20°C. Prior to probing for vRNA, cells were rehydrated, incubated in 0.1% Tween-20 in PBS for 10 min, and mounted on slides. Probing was performed using RNAScope probes and reagents (Advanced Cell Diagnostics) as described before (9, 15). Briefly, coverslips were treated with protease solution for 15 min in a humidified HybEZ oven (Advanced Cell Diagnostics) at 40 °C. The coverslips were then washed with PBS, and pre-designed anti-sense probes specific for SARS-CoV-2 positive strand S gene encoding the spike protein (RNAscope Probe-V-nCoV2019-S, cat# 848561) were applied and allowed to hybridize with the samples in a humidified HybEZ oven at 40 °C for 2 h. The probes were visualized by hybridizing with preamplifiers, amplifiers, and finally, a fluorescent label. First, pre-amplifier 1 (Amp 1-FL) was hybridized to its cognate probe for 30 min in a humidified HybEZ oven at 40 °C. Samples were then subsequently incubated with Amp 1-FL, Amp 2-FL, and Amp 3-FL for 30 min, 30 min, and 15 min, respectively. Between amplifiers, the coverslips were washed with wash buffer (Advanced Cell Diagnostics). After probing for vRNA, nuclei were stained with DAPI (Advanced Cell Diagnostics) at room temperature for 1 min. Finally, coverslips were washed in PBS and then mounted on slides using Prolong Gold Antifade. Images were taken using a Zeiss LSM 880 Airyscan confocal microscope equipped with a ×63/1.4 oil-immersion objective using the Airyscan super-resolution mode. Images were taken of the samples using either the ×63 or ×10 objective. Numbers of nuclei and vRNA puncta in images were quantitated using Volocity 6.3 (PerkinElmer).

### Blockage of SARS-CoV-2 S^E484D^ spread by neutralizing antibodies

SARS-CoV-2 Spike–specific broadly neutralizing antibodies S309, S2X324 and S2K146, which recognize conserved epitopes within the RBD (16) and S2X303, which recognizes the N-terminal domain (NTD) of S (17) were evaluated for their ability to neutralize the SARS-CoV-2 S^E484D^ variant. Cells were then inoculated with SARS-CoV-2 S^E484D^ at an MOI of 1, and virus was removed after 2 h post-infection. Fresh medium containing the corresponding antibodies (10 µg/mL) was added, and cells were harvested at 48 hpi for viral RNA quantification by TaqMan RT-qPCR.

### Antibody internalization assay

Cells were seeded on poly-L-lysine/collagen-coated glass coverslips in 24-well plates at 1× 105 cells per well and allowed to adhere overnight. To prevent non-specific antibody binding through Fc receptors, cells were pre-incubated with Fc receptor blocking reagent (Human TruStain FcX™, BioLegend, catalog no. 422301) at 5 µL per 10^5^ cells for 30 min at 37°C. Cells were then incubated with anti-HA antibody (Alexa Fluor^®^ 594 anti-HA, catlog no. 901511) for H1299 and HCC827 CD71_HA_TMEM106B CTD/CD71_TMEM106B CTD_HA transduced cells, or anti-TMEM106B antibody (Cell Signaling^®^, rabbit monoclonal antibody, catalog no. 93334S) for H522 cells at 500 ng/mL in 1% BSA for 2 h at 37°C to allow antibody binding and internalization. Then cells were washed with ice-cold PBS. Cells were then fixed in 4% paraformaldehyde for 15 min at room temperature. Fixed cells were permeabilized with 0.1% Triton X-100 in PBS for 10 min and blocked with 1% BSA in PBS for 1 h at room temperature. Because the anti-HA antibody was directly conjugated to Alexa Fluor 594, internalized HA-tagged protein was detected without a secondary antibody. For H522 cells, internalized anti-TMEM106B was detected with Alexa Fluor 563-conjugated anti-rabbit IgG, 1:2000 in 1% BSA for 1 h. Lysosomes were labeled with anti-LAMP1 primary antibody (BioLegend, mouse IgG1, catalog no. 328602) followed by Alexa Fluor 488-conjugated anti-mouse IgG, 1:2000 in 1% BSA for 1 h. Nuclei were counterstained with DAPI (H1299 and HCC827) or Hoechst 33342 (H522) 1:2000 1% BSA for 10 min. Coverslips were mounted using Prolong Gold Antifade. Confocal microscopies were taken as previously described using Zeiss LSM 880 Airyscan confocal microscope equipped with a ×63/1.4 oil-immersion objective using the Airyscan super-resolution mode. Colocalization was quantified in Fiji (ImageJ2 version:2.16.0). Multichannel images were split into individual channels. Colocalization between the LAMP1 (green) and HA or TMEM106B (red) channels was assessed by Pearson’s correlation coefficient (R) using the Coloc 2 plugin with automatic thresholding. Pearson’s R values range from −1 to +1, with higher values indicating greater overlap of fluorescence intensities.

### Gradient purification of recombinant viruses

Chimeric VSVs were grown on Vero-TMPRSS2 cells at an MOI of 0.5 in infection media. Cell supernatants were collected upon extensive cytopathic effect and clarified by centrifugation at 1,000 x g for 7.5 min. Supernatants were concentrated using an L-100 XP ultracentrifuge (Beckman Optima) at 22,800 rpm for 90 min in a 70Ti fixed-angle rotor. Pellets were resuspended in 100 mM NaCl, 10 mM Tris pH 7.4, 1 mM EDTA (NTE) at 4°C overnight, and resuspended virus preparations were banded on a 15-45% sucrose-NTE gradient at 35,000 rpm for 3 h in a SW-41Ti swinging-bucket rotor. Virus preparations were extracted by side puncture of tubes, centrifuged again using an L-100 XP ultracentrifuge (Beckman Optima) at 22,800 rpm for 90 min in a 70Ti fixed-angle rotor, and resuspended in NTE at 4°C overnight. Protein content of virus preparations was measured by Bradford Assay (MilliporeSigma) according to the manufacturer’s instructions. Purity of virus preparations was assessed by SDS-PAGE on a 4-15% Tris-glycine gel (Bio-Rad) under reducing and denaturing conditions.

### Biolayer interferometry (BLI)

BLI assays were performed with an Octet RED96e (Sartorius) and analyzed using ForteBio Data Analysis 11.1 software. All reagents were diluted in kinetic buffer (25 mM HEPES pH 8.0, 150 mM NaCl, 1% BSA (w/v)). Briefly, Octet® Anti-Mouse Fc Capture Biosensors (Sartorius) were equilibrated in kinetic buffer for 360 sec and then used to capture anti-SARS-CoV-2 S2 mouse antibody (gift from Diamond lab, made in house) at 50 nM (based on the molecular weight of the dimer) for 300 sec. Biosensors were subsequently dipped into kinetic buffer for 30 sec and then into gradient-purified VSV-SARS-CoV-2 or VSV-SARS-CoV-2-E484D particles at 20 µg/mL for 1800 sec. Biosensors were again dipped into kinetic buffer for 30 sec, then into TMEM106B-Fc at 50 nM (based on the molecular weight of the dimer) for 1800 sec, and finally into kinetic buffer for 300 sec. Baseline measurements were subtracted and sensorgrams are shown.

### Syncytia formation assays

To assess S-mediated cell–cell fusion, Vero-TMPRSS2 cells were seeded in 24-well plates one day prior to infection and inoculated with VSV-GFP-SARS-CoV-2 bearing S-WT, S-E484D or S-E484D/V951A at an MOI of 0.1. H522-TMEM106B-ΔNTD and H661-TMEM106B-ΔNTD cells were inoculated in parallel with VSV-GFP-SARS-CoV-2 bearing S-E484D, S-E484D/V951A, S-E484A, S-E484G or S-E484K at an MOI of 0.1. At 24 hpi (Vero-TMPRSS2) or 48 hpi (H522 and H661 derivatives), cells were fixed with 4% PFA and GFP-positive syncytia were imaged on a Leica fluorescence microscope using a 4× objective. Syncytia were defined as GFP-positive multinucleated cells.

### Bioinformatic analysis of the codon 484 substitution landscape

Spike residue 484 corresponds to nucleotides 23012–23014 of MN908947.3 (NC_045512.2). The parental codon is GAA (Glu). For each of the three positions, all single-nucleotide substitutions were enumerated and the amino acid each produce was assigned using the standard genetic code. Six substitutions are reachable in one nucleotide change: E484A (23013 A>C), E484K (23012 G>A), E484Q (23012 G>C), E484D (23014 A>C or A>T), E484G (23013 A>G) and E484V (23013 A>T). Codon 484 variants were counted in GenBank INSDC genomes through LAPIS/CoV-Spectrum on October 12, 2026. The dataset comprised 9,438,404 genomes. For each amino-acid change at codon 484, the number of genomes carrying it was recorded (E484A 4,306,700; E484K 370,685; E484Q 8,237; E484D 229; E484G 170; E484V not observed). Frequency was calculated as count divided by 9,438,404. Lineage annotations are those assigned in the database: E484A (Omicron-defining), E484K (Beta, Gamma and Mu), E484Q (Kappa). E484D and E484G had no lineage assignment. Figure 10A was drawn in R 4.3.1 with ggplot2 3.4.4 and scales 1.2.1. Frequencies are plotted on a logarithmic axis. Changed bases are colored by lineage, and E484D is shown in teal. Plotting scripts were written with the assistance of Claude Code (Anthropic, version 2.1.309).

SARS-CoV-2 sequencing runs with an E484D call (A>C or A>T at nucleotide 23014) were retrieved from NCBI Sequence Read Archive on October 12, 2026, using the variant calls reported by NCBI for each run. 406 runs carried an E484D call. For each run, the allele fraction was calculated as alternate read depth divided by total read depth at position 23014, using the depths in the call table. Each run contributes one point. Each run was assigned one context from its SRA metadata (host, isolation source, study title and BioProject), in the following order. (1) Wastewater: sample type or title indicates wastewater. (2) Cell culture: isolation source includes Vero, cell culture, HEK, cell line, or supernatant, host is Chlorocebus aethiops, or the BioProject is a laboratory cell-culture study. (3) Animal: host is hamster or Mus musculus, or the title includes hamster, mouse adapted, or serial passag. (4) Wild animal: host is Odocoileus virginianus (free-ranging deer). (5) Drug-selection passages: BioProjects PRJNA1035781 (molnupiravir resistance selection, 17 runs; SARS-CoV-2 passaged in Vero E6 cells with NHC, MRK-A or no drug, each of 30 passages was sequenced by whole-genome sequencing.) and PRJNA692078 (remdesivir resistance evolution, 6 runs). (6) Clinical surveillance: host is Homo sapiens, or the source includes swab, clinical, patient, nasal, nasopharyngeal or sputum; the single run from the monoclonal-antibody treatment BioProject PRJEB55794 is classed here. (7) Unspecified: none of the above. In vivo: clinical surveillance (n = 227, including the monoclonal-antibody run) and animal (n = 17, combining wild deer and experimental animals). Ex vivo: cell culture, including drug-selection passages (n = 39), and wastewater (n = 14). Context not recorded: unspecified (n = 111). Horizontal bars show the median allele fraction in each column. The MA1 mouse-adaptation values (E484D at 96% of reads in mouse lung and 92% in cell supernatant) were taken from the supplementary data of the MA1 study and are plotted as open diamonds, since they come from a different study and sequencing method.

### Bioinformatic analysis of brain and lung E484A frequencies in a mouse model

Minority variants at spike codon E484 (genome position 23012-23014, MN908947.3) were identified from the sequencing libraries and alignments described in (22); no additional sequencing was performed for this analysis. Briefly, cDNA was generated with random hexamers, viral genomes amplified with ARTIC v4 primers, libraries prepared with the SeqWell plexWell 384 kit and sequenced on an Illumina MiSeq (V2, 500 cycles). Reads were adaptor-and quality-trimmed with Trimmomatic v0.36, aligned to the SARS-CoV-2 reference genome (MN908947.3) with bwa v0.7.15, and pile-ups generated with samtools v1.9. Per-position nucleotide counts and frequencies were taken from the iVar v1.2.2 variant output generated with a minimum depth of 10, a minimum base quality score of 20 and a consensus frequency threshold of 0 (that is, majority base as consensus), so that all sub-consensus alleles passing the quality filter were retained. For this sub-analysis, frequency estimates were restricted to positions covered by at least 100 reads, consistent with the depth threshold used for the Shannon entropy calculations, to ensure robustness of the measurement; samples failing this threshold at the codon of interest were excluded from the analysis rather than recorded as zero. The frequency of the E484A-encoding substitution was extracted for each brain and lung sample and expressed as the proportion of reads carrying the alternative allele at that position. Samples that met the depth criterion but in which no E484A-supporting reads were called by iVar were assigned a frequency of 0 and are plotted as such. The sequenced viral inoculum of the WA reference genome was processed through the identical pipeline and served as the baseline for both background sequencing error and pre-existing variation in the stock.

### Statistical analysis

Statistical analyses were performed in GraphPad Prism. Unless otherwise indicated in the figure legends, data are presented as the mean of three independent replicates with error bars showing the standard error of the mean (SEM). Comparisons between two groups were made with a two-tailed unpaired Student’s t test, and comparisons of three or more groups against a single control were made with a one-way ANOVA with Dunnett’s correction for multiple comparisons. Viral RNA copy numbers were log-transformed prior to statistical testing. P values < 0.05 were considered statistically significant; ns, not significant; *, P < 0.05; **, P < 0.01; ***, P < 0.001; ****, P < 0.0001.

## ACKNOWLEDGEMENTS

This study was supported by the Sondra Schlesinger Fellowship to M.X., NIH T32 training grant (T32CA009547-34) to K.M.L., NIH AI157155 to M.S.D. and NIH AI163019 to S.P.J.W. We thank all members of the Kutluay and Diamond laboratories for their critical feedback and input during the course of this project.

